# Pathological turret mutations in the cardiac sodium channel cause long-range pore disruption

**DOI:** 10.1101/2021.01.15.426807

**Authors:** Zaki F Habib, Manas Kohli, Samantha C Salvage, Taufiq Rahman, Christopher L-H Huang, Antony P Jackson

**Author notes:** **Authorship note:** ZFH and MK made an equal contribution to this work. Address correspondence to: Antony P Jackson, Department of Biochemistry, University of Cambridge, Cambridge, UK CB2 1QW, UK.; Christopher L-H Huang, Department of Physiology, Development and Neuroscience, University of Cambridge, CB2 3EG, UK.; Zaki F Habib, Department of Physiology, Development and Neuroscience and Department of Biochemistry, University of Cambridge, UK.

## Abstract

The voltage-gated sodium channel Nav1.5 initiates the cardiac action potential. Germline mutations that disrupt Nav1.5 activity predispose affected individuals to inherited cardiopathologies. Some of these Nav1.5 mutations alter amino acids in extracellular turret domains DII and DIII. Yet the mechanism is unclear. In the rat Nav1.5 structure determined by cryogenic electron microscopy, the wild-type residues corresponding to these mutants form a complex salt-bridge between the DII and DIII turret interface. Furthermore, adjacent aromatic residues form cation-π interactions with the complex salt-bridge. Here, we examine this region using site-directed mutagenesis, electrophysiology and *in silico* modeling. We confirm functional roles for the salt-bridges and the aromatic residues. We show that their disruption perturbs the geometry of both the DEKA selectivity ring and the inner pore vestibule that are crucial for sodium ion permeability. Our findings provide insights into a class of pathological mutations occurring not only in Nav1.5 but also in other sodium channel isoforms too. Our work illustrates how the sodium channel structures now being reported can be used to formulate and guide novel functional hypotheses.

## Introduction

The voltage-gated sodium channel isoform Nav1.5 (the product of the *SCN5A* gene) is mainly expressed in cardiac muscle, where it initiates the depolarizing phase of the action potential (1). The structure of rat Nav1.5 has been determined by cryogenic electron microscopy (cryo-EM) to a resolution of 3.2-3.5Å (2). This single, 250 kDa protein contains four homologous domains (DI-DIV), each of which contains six transmembrane α-helices (S1-S6), arranged with four-fold rotational pseudosymmetry. In each domain, helices S1-S4 form an outer voltage-sensing segment. Helices S5 and S6, together with re-entrant P loop helices, P1 and P2, form the central ion-conducting pore (Figure 1A and B).

**Figure 1.**
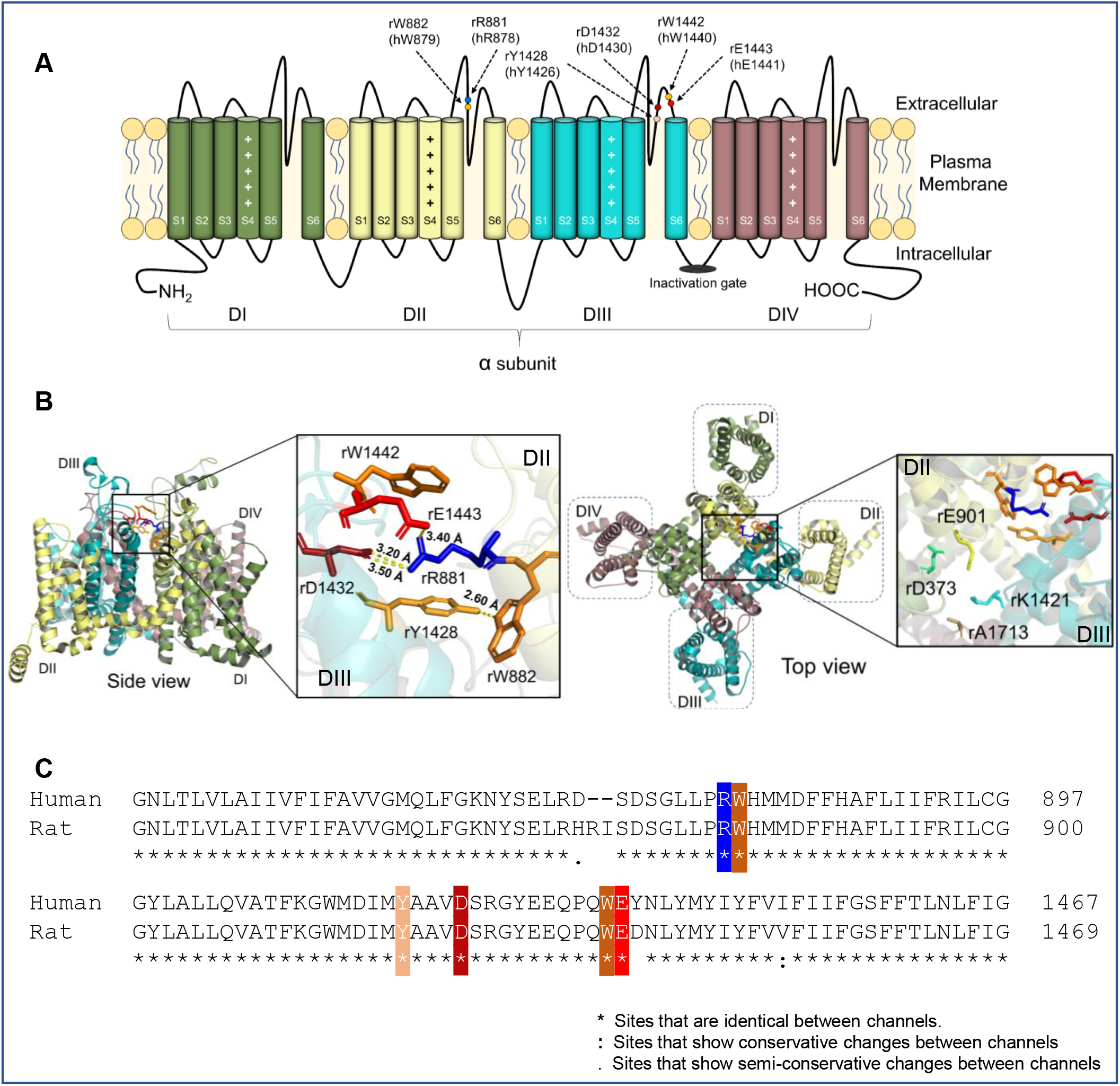
A complex salt bridge and cation-π interactions within the Nav1.5 DII-DIII extracellular turret. **(A)** Schematic representation of the Nav1.5 channel. The four homologous domains, DI-DIV are indicated. The helices S5 and S6 line the channel pore and the positively-charged S4 helices function as the voltage sensor. The positions of amino acid residues at the extracellular turret DII-DIII interface studied in this paper are indicated for both human (h) and rat (r) Nav1.5 sequences. **(B)** A cartoon representation of the rat Na_v_1.5 structure (PDB ID: 6UZ3), showing the topology of the channel from the side and from above. With the DII-DIII turret interface residues and the DEKA selectivity ring highlighted. Note the different domain orientations in the two expanded inserts. Distances between amino acids are indicated in Å. **(C)** primary sequence alignments of human Nav1.5 (Uniprot ID: Q14524-1) and rat Nav1.5 (Uniprot ID: P15389-1). Residues discussed in the text are highlighted.

Each S4 helix contains a series of positively charged amino acids that sense the membrane potential, such that they are displaced in the extracellular direction following membrane depolarization. This movement induces a series of conformational changes In domains DI, DII and DIII, that leads to the widening of the inner pore between the intracellular faces of the S6 helices and thus initiates channel activation (3). However, movement of the DIV S4 helix - which occurs a few milliseconds after channel opening - facilitates binding of the intracellular fast inactivation gate (Figure 1A) to a site between the intracellular faces of the DIV, S5 and S6 helices. This binding drives the channel into an inactivated state. As the resting potential is restored, all four S4 helices return to their original positions and the inactivated channel is reset into the closed state, where it can again respond to further depolarization signals (4).

On the extracellular face of the pore, the re-entrant P1 and P2 helices from each domain assemble into an inverted and truncated cone-like structure. At its narrowest point (the constriction site), lie the highly conserved residues: Asp (DI); Glu (DII); Lys (DIII) and Ala (DIV). These form the DEKA selectivity ring, whose sequence and geometry is critical for selective sodium ion permeability (5). In the rat Nav1.5 channel, the selectivity ring residues are D373; E901; K1421 and A1713 (Figure 1B). Within each domain, the extracellular loop regions between the S5 helix and P1 and between P2 and the S6 helix extend outward, such that they form an exterior turret surrounding and shielding the outer pore vestibule (Figure 1 A and B) (6).

Germline *SCN5A* mutations that reduce or abolish channel activity are associated with inherited, autosomal dominant cardiopathologies, including Brugada syndrome (BrS) and sinus node dysfunction (SND). BrS is characterized by familial atrial fibrillation, ventricular tachyarrhythmias and syncope. SND is also associated with tachyarrhythmias involving the sinus node. Although in practice, the BrS and SND phenotypes can clinically overlap. Together, these syndromes account for a significant fraction of sudden death in young to middle-aged patients (7). Because almost all BrS and SND patients are heterozygous, they typically exhibit about a 50% reduction in the level of functional Nav1.5 compared to phenotypically normal individuals. This reduction can compromise action potential conduction velocities and precipitate arrhythmogenicity (8).

Several *SCN5A* deletion and missense mutations associated with BrS and SND, prevent the folding of the Nav1.5 channel and lead to its retention in the endoplasmic reticulum (ER) (9). Other missense mutations do not prevent folding but disrupt channel activity because they occur in regions known to be functionally important, such as the S4 helices or the inactivation gate (10). However, some BrS/SND missense mutations occur in regions whose association with sodium channel function is less clear. Such mutations are likely to be particularly informative, both structurally and therapeutically, as they may reveal new insights into Nav channel behavior. This class of mutation is exemplified in an individual with a BrS/SND-like pathology who was heterozygous for the Nav1.5 missense mutation R878C, a residue located on the DII extracellular turret loop connecting the S5 helix and P1. When tested by heterologous expression, the mutant displayed a complete loss of function, despite normal Nav1.5 expression on the plasma membrane (11). Two similar BrS-associated Nav1.5 mutations, D1430N and E1441Q are located on the DIII extracellular turret loop connecting P2 to the S6 helix (Figure 1 A and B). The D1430N mutant displays a complete loss of function (12). In the case of E1441Q, no experimental analysis of its gating behavior has yet been described (13). The human R878, D1430 and E1441 residues and their immediate neighbours are conserved in the rat Nav1.5 sequence (Figure 1C). Thus, by using the rat structure as a guide, the functional significance of these pathological mutations can now be examined in their broader structural context. It should be noted, however, that there are small insertions and deletions between the two sequences, so the corresponding residue positions are not identical. Thus, human R878 is equivalent to rat R881, human D1430 corresponds to rat D1432 and human E1441 corresponds to rat E1443. For clarity, both the human and the equivalent rat Nav1.5 residues that we have experimentally studied are indicated in Figure 1A and are summarized in Table 1.

**Table 1.** Summary tables. Left: The equivalent human (h) and rat (r) residues for the amino acids studied in this current work are listed. Right: The human (h) Nav1.5 mutations used in the electrophysiological and biochemical assays and the equivalent rat (r) Nav1.5 mutants examined by MD simulation are shown.

| Equivalent human and rat Nav1.5 residues |  | Human Nav1.5 mutations used in functional assays | Equivalent rat Nav1.5 mutations modeled by MD simulation |
| --- | --- | --- | --- |
| Human | Rat |  |  |
| R878 | R881 | W1440Y | W1442Y |
| W879 | W882 | W1440A | — |
| Y1426 | Y1428 | Y1426F | — |
| D1430 | D1432 | Y1426W | — |
| W1440 | W1442 | Y1426A | — |
| E1441 | E1443 | E1441D | E1443D |
|  |  | D1430E | — |
|  |  | E1441Q | E1443Q |
|  |  | R878C | — |
|  |  | W879Y | - |
|  |  | W879A | - |
|  |  | R878E/E1441R | — |
|  |  | R878C/E1441C | — |
|  |  | R878D/D1430R | R881D/D1432R |
|  |  | R878C/D1430C | - |

Interestingly, in the rat Nav1.5 structure, the positively charged guanidinium group of R881 on the DII turret lies less than 4Å from the negatively charged carboxylate groups of D1432 and E1443 on the adjacent DIII turret (Figure 1B) (2). This arrangement is consistent with a complex salt-bridge, in which more than two charged residues interact (14, 15). Moreover, the charged residues are sandwiched between the side chains of rat Y1428 (equivalent to human Y1426) and rat W1442 (equivalent to human W1440) on the DIII turret, such that both the aromatic rings lie in register with, and less than 4Å from, the delocalised positive charge on the R881 guanidinium group. This is consistent with cation-π interactions (16). In the rat structure, residue Y1428 on the DIII turret also forms a hydrogen bond with W882 (equivalent to human W879) on the DII turret (Figure 1B). It is reasonable to assume that the corresponding residues in human Nav1.5 adopt the same structural relationships as they do in rat Nav1.5; that they act together to stabilise the DII, DIII turret interface and that the contacts are required for Nav1.5 function.

Here we show that mutational changes to these residues in human Nav1.5 abolish or modify channel activity, yet do not affect surface expression of the mutant channels. Using molecular dynamics (MD) simulations based on the rat Nav1.5 structure (2), we show that the disabling mutations alter the geometry and stability of the DII-DIII turret interface loop and induce longer-range destabilizing changes to the geometry of the DEKA selectivity ring and the inner pore vestibule. Consequently, these changes are likely to prevent the passage of sodium ions through the pore.

## Results

### Electrophysiological properties of the wild-type and mutant Nav1.5 channels

A summary of the human Nav1.5 mutants generated for the electrophysiological experiments is presented in Table 1. We first examined the human pathological BrS/SND mutation, R878C (11) and the putative human BrS mutation, E1441Q (13). In agreement with previous work (11), we failed to detect any electrophysiological activity from the R878C mutant (Figure 2A). The E1441Q mutation was similarly inactive, thus corroborating its previously inferred role as causative for BrS (13) (Figure 2A). Given the structural insights noted above, this data supports a functional role for a salt-bridge between human R878 and E1441. Previous work had established that the conservative, human Nav1.5 mutant R878K was also completely inactive (11), suggesting strict structural constraints on the salt-bridge. To extend this observation and guided by the rat Nav1.5 structure, we generated individual conservative mutations in human Nav1.5: E1441D and D1430E and tested them for electrophysiological behavior compared to wild-type human Nav1.5. The E1441D mutant displayed a 93% reduction in peak current activity, compared to the wild-type value. Nevertheless, the currents were large enough to measure additional gating parameters. Compared to wild-type Nav1.5, the E1441D mutant displayed a significantly increased slope factor (*k*) for activation. However, there was no effect of the E1441D mutation on the steady-state V_1/2_ of activation or inactivation or the slope factor (*k*) for inactivation. By contrast, there was no detectable current from the D1430E mutation (Figure 2A-C; Figure 3A and B; Table 2).

**Table 2.** Parameters from whole-cell patch clamp steady-state activation and inactivation data. Both activation and inactivation data were fit using a Boltzmann function to deduce all the parameters. The data are expressed as mean ± SEM and One-Way ANOVA statistical analysis followed by Dunnett post-hoc tests performed to determine the statistical significance between each mutant and the wild-type Nav1.5 channel. *p<0.05, **p<0.01, ***p<0.001, ****p<0.0001.

| Activation |  |  |  |  | Inactivation |  |  |
| --- | --- | --- | --- | --- | --- | --- | --- |
| HEK293F cells | Peak $I_{Na}$ | $V_{1/2}$(mV) | k(mV) | n | $V_{1/2}$(mV) | k(mV) | n |
| WT | -332.29 ± 36.29 | -46.89 ± 2.43 | 5.22 ± 0.35 | 9 | -84.75 ± 1.702 | -7.925 ± 0.235 | 14 |
| hW1440Y | -342.46 ± 42.15 | -47.89 ± 1.47 | 5.27 ± 0.45 | 8 | -82.52 ± 2.593 | -7.351 ± 0.326 | 9 |
| hW1440A | -60.85 ± 13.00**** | -41.77 ± 1.51 | 7.85 ± 0.40*** | 9 | -80.49 ± 3.272 | -7.606 ± 0.455 | 9 |
| hY1426F | -180.47 ± 40.13** | -44.4 ± 0.95 | 5.98 ± 0.28 | 8 | -85.57 ± 1.958 | -7.894 ± 0.238 | 9 |
| hY1426W | -44.23 ± 7.54**** | -41.83 ± 1.84 | 7.86 ± 0.47*** | 8 | -78.48 ± 3.273 | -7.942 ± 0.364 | 8 |
| hY1426A | -68.44 ± 11.25**** | -41.61 ± 1.41 | 7.22 ± 0.17** | 7 | -79.81 ± 3.474 | -7.888 ± 0.461 | 8 |
| hE1441D | -24.83 ± 2.23**** | -40.57 ± 1.55 | 8.16 ± 0.64**** | 7 | -82.17 ± 2.4 | -7.394 ± 0.457 | 12 |

**Figure 2.**
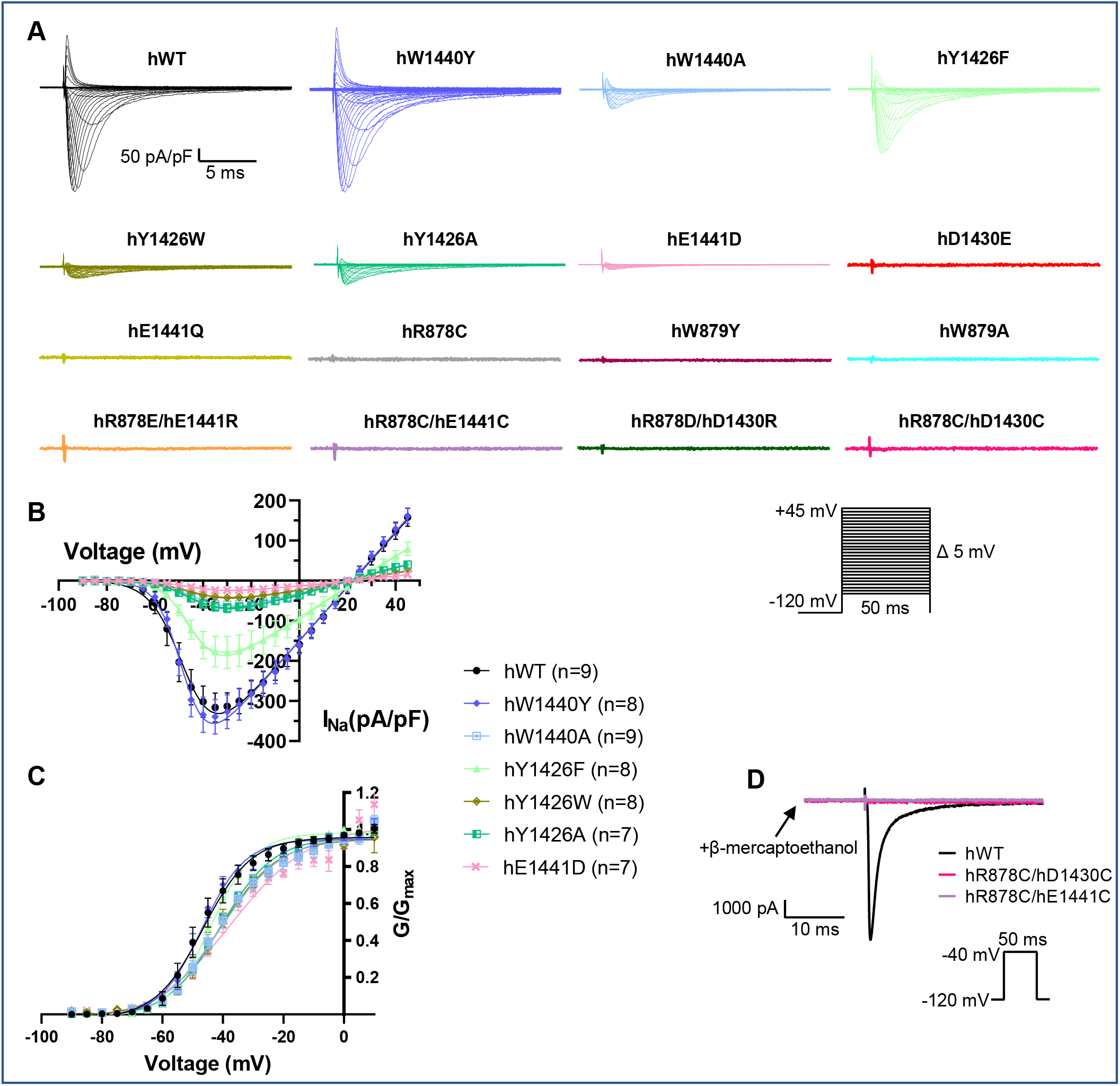
Steady-state activation properties of wild-type and mutant human Nav1.5 channels. **(A)** Typical whole-cell current traces from HEK293F cells transfected with human (h), wild-type or mutant Nav1.5 channel under the activation protocol as outlined in Methods. **(B)** Currents, normalized to whole-cell capacitance, versus voltage curves for the wild-type and active mutants. **(C)** Channel conductance as a function of voltage for wild-type and active mutants. *Curves* were fit to a Boltzmann function as outlined in Methods. The data are means ± SEMs. (*n* ≥ 7). Statistical significance was tested with one-way ANOVA. Compared to wild-type channels, no differences were observed in V_1/2_(activation). However, the slope factor (*k*), of human: W1440A, Y1426W, Y1426A and E1441D were significantly different from the wild-type value. See **Table 2** for individual values. **(D)** The disulfide reducing agent β-mercaptoethanol does not rescue human: R878C/E1441C or the human: R878C/D1430C double mutants.

**Figure 3.**
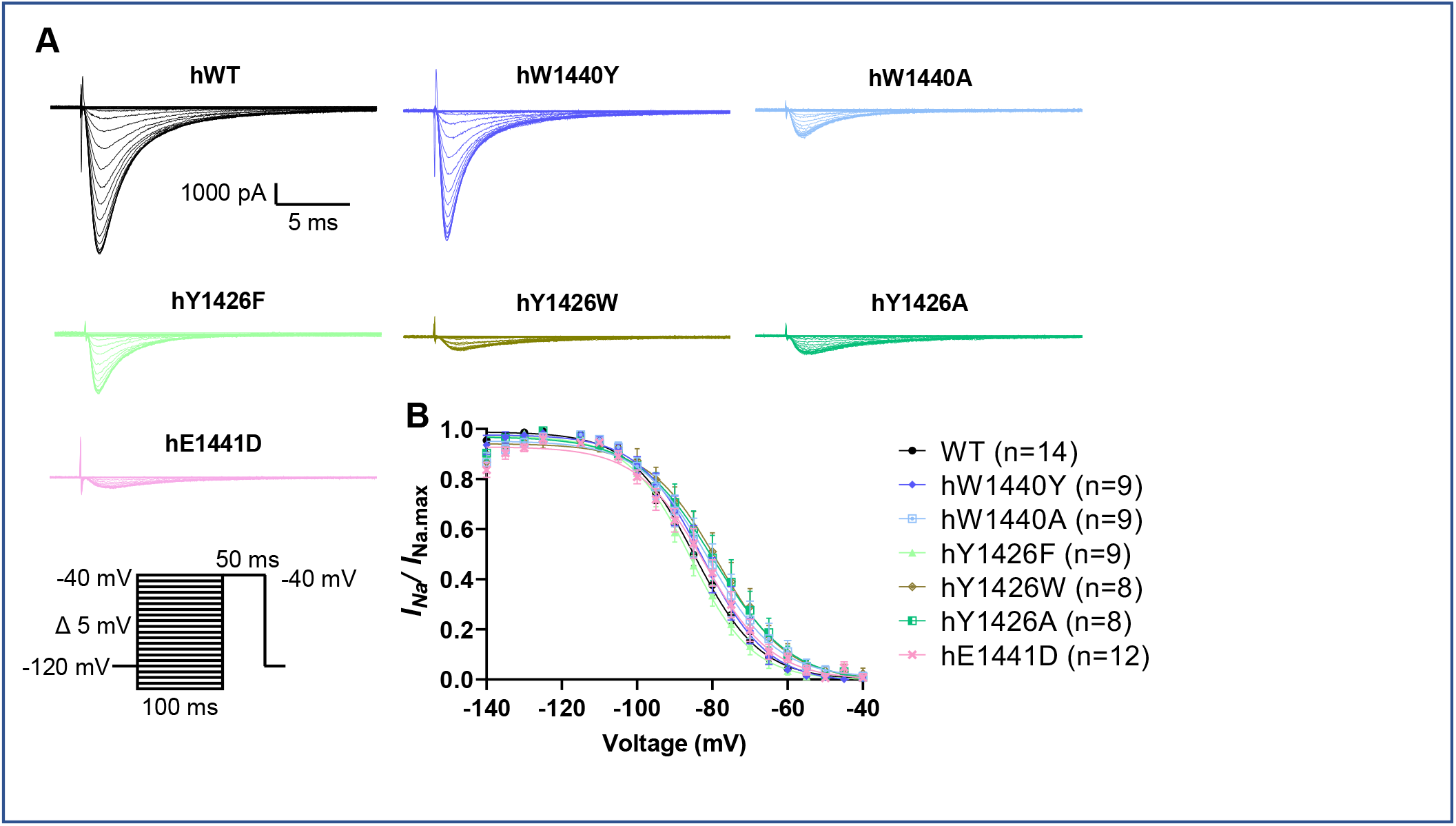
Steady-state inactivation properties of wild-type and the functional mutant human Nav1.5 channels. **(A)** Typical whole-cell current traces from HEK293F cells transfected with human (h), wild-type or mutant Nav1.5 channel under the steady-state inactivation protocols described in Materials. Only those mutants that produced detectable current traces in **Figure 2A** were examined. **(B)** Current-voltage curve with peak I_Na_ of each step normalized with maximum peak I_Na_ and plotted against test pre-pulse potential. The data are means ± SEMs (n ≥ 8, See **Table 2** for individual values) and statistical significance was tested with one-way ANOVA. No shifts in the steady-state inactivation properties were observed in the mutants when compared to the wild-type channel (see **Table 2** for individual values).

A charge-swap approach has been previously used to study ion channel salt bridges (17, 18). Here, the positively and negatively charged residues are reversed by site-directed mutagenesis. The extent to which a given function is maintained or modified by this procedure reflects the structural constraints on the salt bridge in the wild-type protein. We generated a double mutation in human Nav1.5: R878E with E1441R and a separate double mutation: R878D with D1430R. Strikingly, neither of the charge swap mutants exhibited any detectable currents (Figure 2).

As a complementary approach, we also used cysteine-substitution. This method has provided important structural and functional insights into ion channel behavior (19, 20). Here, the D1430 and E1441 residues of the human R878C mutant were separately replaced with cysteine. We expected that if any activity was restored in these mutants, it would indicate that a disulfide bond could form between the two DII and DIII turret interface cysteines and provide a functional replacement for the salt bridges (20). However, neither mutation (either with or without prior reduction) exhibited any detectable current (Figure 2D). Taken together, these experiments emphasize the tight functional constraints that apply to the complex salt bridge residues (see Discussion).

We next examined the functional importance of the aromatic residues surrounding the complex salt bridge (Figure 1B). The human W1440Y mutation was functional and showed no statistically significant differences in peak current, or steady-state gating parameters compared to human wild-type Nav1.5 In contrast, the human W1440A mutation, which lacked any aromatic residue at this position, had a significantly (~80%) reduced peak current relative to wild-type human Nav1.5 and a significantly increased slope factor for activation. Again, however, the W1440A mutation had no significant effect on steady-state gating parameters for inactivation (Figure 2A-C; Figure 3A and B; Table 2). The functional importance of the Y1426 aromatic residue in human Nav1.5 was similarly investigated by replacing it with alanine. The Y1426A mutant displayed a 70% reduction in peak current relative to wild-type Nav1.5 and a significant increase in (*k*) activation, but with no significant effect on steady-state gating parameters (Figure 2A-C; Figure 3A and B; Table 2). Thus, the separate removal of each of the aromatic groups that form the putative cation-π interactions with the R878 guanidinium group, significantly reduced but did not abolish channel activity.

Based on the rat Nav1.5 structure, we infer that human Y1426 on the DIII turret additionally forms a hydrogen bond with human W879 on the DII turret (Figure 1B). To test the importance of this interaction, we first replaced the human Y1426 residue with a bulkier tryptophan residue. This change would be expected both to abolish the hydrogen bond with human W879 and to also perturb the packing at the DII and DIII turret interface, whilst retaining the cation-π bond itself. The human Y1426W mutation displayed a significant reduction in peak current and an increase in the slope factor (*k*) for activation (Figure 2A-C; Figure 3A and B; Table 2). To specifically test the role of the hydrogen bond, we generated a human Y1426F mutant. This mutant should largely retain its cation-π bonding potential relative to wild-type human Nav1.5, with minimum disturbance to side chain packing, whilst being unable to form a hydrogen bond with W879. The human Y1426F mutant displayed a 50% reduction in peak current, with no significant effect on steady-state gating parameters (Table 2). Finally, we examined the role of the human W879 residue by generating a human W879A and a W879Y mutant. No current responses could be detected in either of these cases (Figure 2A). Since this was a more severe phenotype than displayed by the Y1426F mutant, it suggests an important stabilising role for the tryptophan residue at position 879 of human Nav1.5, acting in addition to its hydrogen bonding potential with Y1426.

In addition to steady-state parameters, we also examined recovery from inactivation kinetics (Figure 4A and B). As noted previously (21), recovery from inactivation was best fitted to a double exponential curve corresponding to a fast and slow component. However, there were no statistically significant differences in the kinetics of either of these components between any of the mutants and wild-type channels (Table 3).

**Table 3.** Parameters form whole-cell patch clamp recovery from inactivation data. A double exponential function was used to fit the data and expressed as mean ± SEM. Statistical significance was analysed using a one-Way ANOVA, followed by Dunnett post-hoc tests performed to determine the statistical significance between each mutant and the wild-type Nav1.5 channel. *p<0.05, **p<0.01, ***p<0.001, ****p<0.0001.

| Recovery from Inactivation |  |  |  |  |  |  |  |
| --- | --- | --- | --- | --- | --- | --- | --- |
| | $k_{fast}$ | $k_{slow}$ | $\tau_{fast}$ (ms) | $\tau_{slow}$ (ms) | $t_{1/2 \text{ fast}}$ (ms) | $t_{1/2 \text{ slow}}$ (ms) | n |
| HEK293F cells |  |  |  |  |  |  |  |
| WT | $0.122 \pm 0.008$ | $0.022 \pm 0.002$ | $8.611 \pm 0.524$ | $48.47 \pm 3.747$ | $5.969 \pm 0.363$ | $33.60 \pm 2.597$ | 12 |
| hW1440Y | $0.183 \pm 0.021$ | $0.029 \pm 0.004$ | $5.905 \pm 0.669$ | $39.05 \pm 5.21$ | $4.093 \pm 0.464$ | $27.07 \pm 3.610$ | 7 |
| hW1440A | $0.144 \pm 0.015$ | $0.026 \pm 0.002$ | $7.472 \pm 0.709$ | $39.99 \pm 3.431$ | $5.179 \pm 0.492$ | $27.72 \pm 2.378$ | 8 |
| hY1426F | $0.120 \pm 0.010$ | $0.020 \pm 0.002$ | $8.658 \pm 0.757$ | $54.45 \pm 5.037$ | $6.000 \pm 0.524$ | $37.74 \pm 3.492$ | 6 |
| hY1426W | $0.185 \pm 0.040$ | $0.026 \pm 0.005$ | $6.709 \pm 1.332$ | $48.08 \pm 10.54$ | $4.650 \pm 0.923$ | $33.33 \pm 7.302$ | 6 |
| hY1426A | $0.152 \pm 0.020$ | $0.035 \pm 0.011$ | $7.377 \pm 1.234$ | $42.64 \pm 10.25$ | $5.113 \pm 0.855$ | $29.55 \pm 7.107$ | 6 |
| hE1441D | $0.160 \pm 0.020$ | $0.022 \pm 0.002$ | $6.999 \pm 1.221$ | $49.04 \pm 4.993$ | $4.852 \pm 0.846$ | $33.99 \pm 3.461$ | 6 |

**Figure 4.**
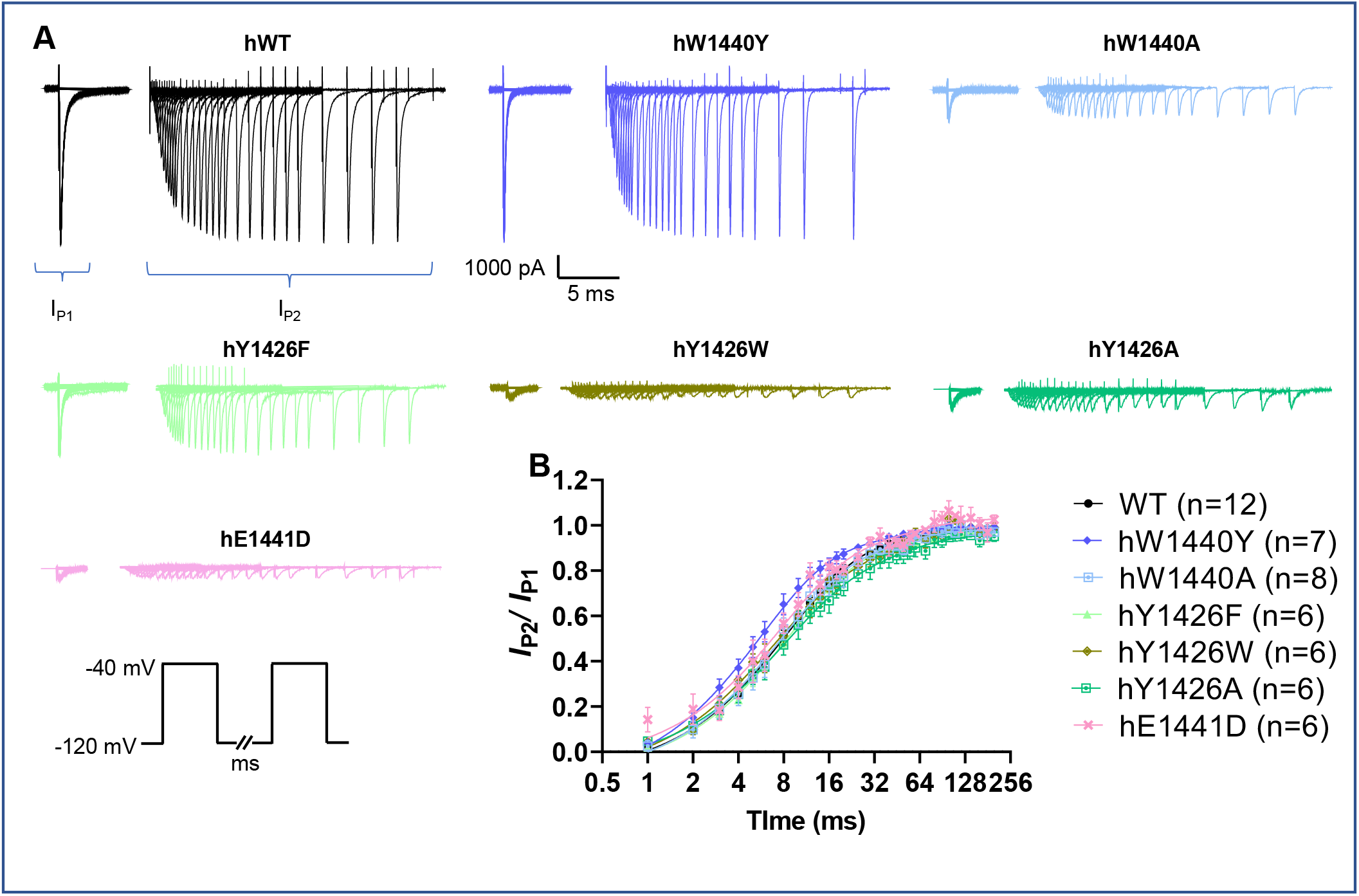
Recovery from inactivation properties of wild-type and the active mutant human Nav1.5 channels. **(A)** Representative whole-cell traces from recovery from the inactivation protocol as outlined in Methods. (**B)** The fraction of human (h), wild-type or mutant Nav1.5 channel I_Na_ recovering from the inactive state plotted against the increasing time duration between the two depolarizing pulses. Only those mutants that produced detectable current traces in **Figure 2A** were examined. No significant differences were observed between the wild-type and mutant channels (see **Table 3** for individual values).

### The Nav1.5 mutants are expressed on the plasma membrane

A major effect of the mutations is on the magnitude of their peak currents relative to wild-type Nav1.5 (Table 2). This could indicate that severely compromised mutants, particularly those mutants that lack detectable gating activity, fail to traffic to the plasma membrane. To test this possibility, we measured the fraction of human Nav1.5 channels on the plasma membrane of the transfected HEK293F cells, using a surface biotinylation approach (22, 23). Both the wild-type human Nav1.5 and all the mutant channels were detected on the plasma membrane and to a similar extent. There was no correlation between surface expression and peak currents for any of the channels (Figure 5A and B). This result agrees with the previous report that the R878C mutant was expressed on the plasma membrane (11). We conclude that the mutations introduce relatively subtle structural changes into the Nav1.5 channel and do not significantly compromise global folding. Yet at the same time, they impair one or more steps in the gating mechanism.

**Figure 5.**
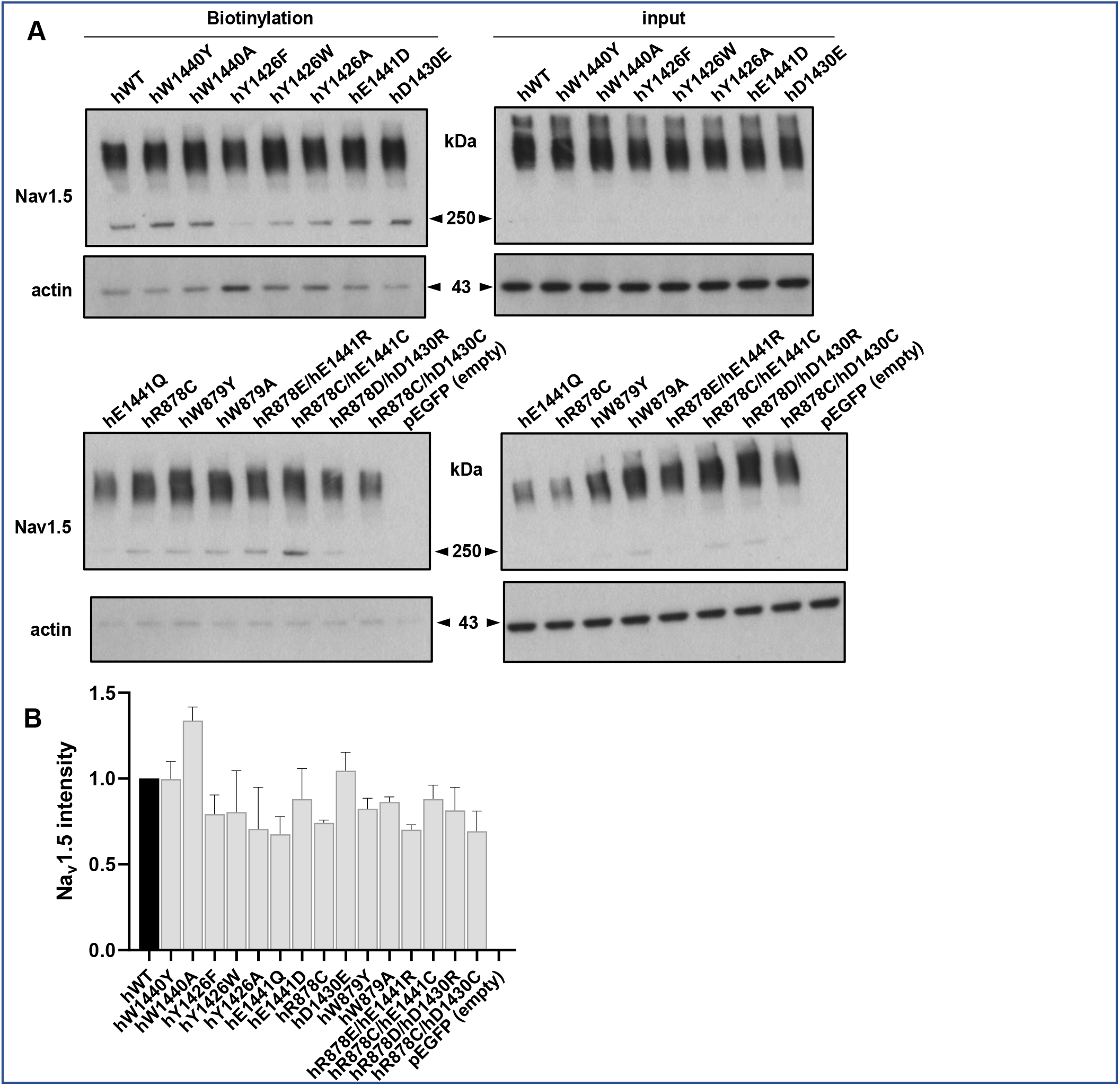
Cell surface biotinylation assay. **(A)** Representative western blots of biotinylated cell surface fraction and total lysate of HEK293F cells transiently transfected with wild-type or mutant human (h), wild-type and mutant Nav1.5 channels. Actin was used as a loading control. (**B)** Quantitation of the Nav1.5 intensities (n=3). Normalized intensities for each channel are presented relative to the normalized intensity for the wild-type Nav1.5. No significant differences were seen between the wild-type and mutant channels.

### Veratridine fails to rescue the human Nav1.5 null-mutations

The steroidal alkaloid Nav channel activator veratridine inhibits channel transitions from the open to the inactivated state (24). Veratridine binds to an intracellular site, located between DI and DIV, that is distinct and quite separate from the extracellular DII-DIII turret interface (25). If the turret null-mutations were driving the channel into the inactivated state, then we would expect veratridine to overcome, or at least attenuate inactivation and hence restore some activity. If on the other hand, the mutations were preventing the earlier activation step, then veratridine should not influence these channels. We found that 100 μM veratridine, a concentration that is known to be saturating for wild-type Nav1.5 (26), induced a clear tail current in the wild-type human Nav1.5 channel, but failed to rescue any of the human Nav1.5 null-mutants (Figure 6A and B). Therefore, the Nav1.5 mutations most likely interfere with conformational movements normally required for the activation step, for example pore opening.

**Figure 6.**
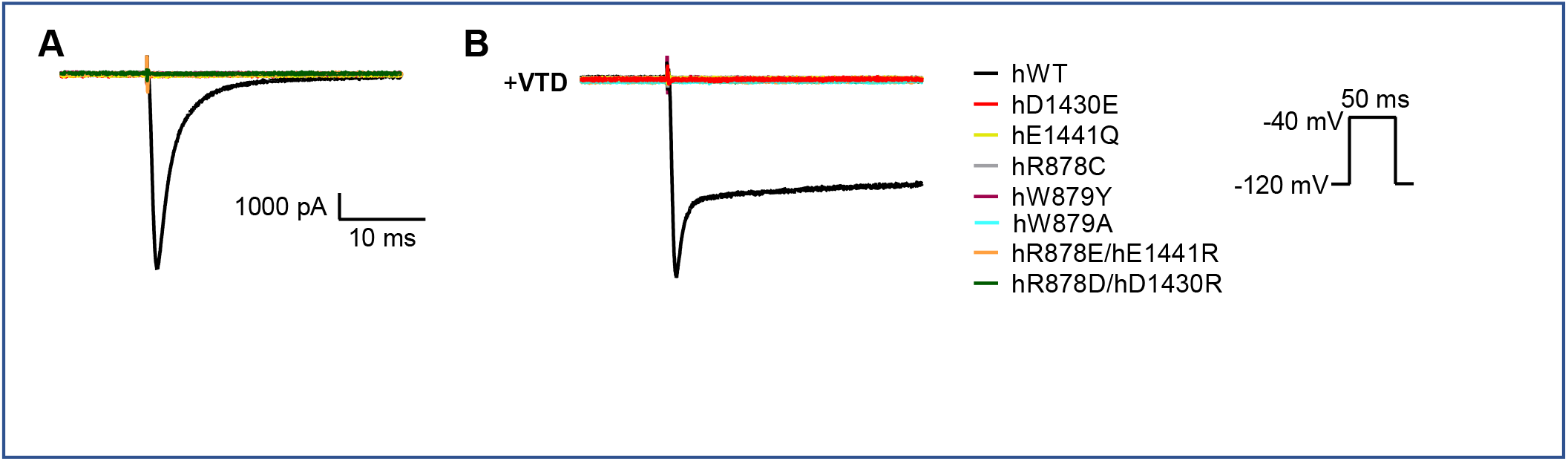
The sodium channel activator veratridine does not rescue human Nav1.5 null-mutants. The gating responses of the human (h), Nav1,5 wild-type (WT) and null-mutants were tested **(A)**, in the absence or **(B)**, in the presence of 0.1 mM veratridine (VTD) (0.1 mM). Representative traces for the wild-type Nav1.5 channel show the classic tail current, caused by veratridine-induced stabilization of the active Nav channel state (26). However, none of the mutants showed any sign of re-activation.

### Atomistic molecular dynamics (MD) simulations of rat Nav1.5 wild-type and mutant structures

Based on the electrophysiological, biochemical and pharmacological evidence presented above, we hypothesized that mutations in the DII-DIII turret interface of Nav1.5 perturb the turret structure, which then allosterically disrupts pore geometry and/or global channel flexibility, to impair sodium ion permeability. To address this question, we carried out atomistic MD simulations (27, 28) of the available wild-type rat Nav1.5 structure and mutants generated by *in silico* mutagenesis (29). Since it was impractical to simulate all the mutants that we characterised experimentally, we chose to model a sub-set of specific mutants, both pathological and artificial, that together represent illustrative examples from the range of recorded gating activities. For these experiments, we used the existing rat Nav1.5 structure as a template (2). The mutations that we examined by MD simulation are summarized in Table 1 and represent three functional classes, as informed by the electrophysiological data (Table 2). They are:

(i) Rat W1422Y. This is the rat equivalent to human W1440Y, a mutation that incorporates a natural Nav channel isoform polymorphism into a Nav1.5 background and whose electrophysiological gating behavior does not differ from the wild-type protein.
(ii) Rat E1443D. This is the rat equivalent to human E1441D, a mutation with a conservative charge replacement, yet with a significantly reduced peak current and altered slope factor.
(iii) Rat E1443Q and rat R881D/D1432R. These mutants are equivalent to the human putative BrS mutant E1441Q and the human R878D/D1430R mutant respectively. Both human mutants produce inactive channels.

For all structures simulated, we assessed channel stability as measured through Root-Mean Square Deviation (RMSD) values. Pore and turret interface stabilities were assessed through RMSD values, energies, and residue distances in each structure.

### Mutations of the turret interface affect both global and local stability of the Nav1.5 structure

The RMSD backbone value for the wild-type structure stabilized at about 7Å for the latter half (5 ns and onwards) of the time course. Hence, we used a 15 ns duration as the minimum length of our comparative simulation work. In our models, we examined the structural flexibility of the whole channels (Figure 7A); the residues of the S5-S6 helices that line the inner pore (Figure 7B) and the residues within the turret interface (Figure 7C). For the case of global stability and the stability of the S5-S6 pore residues, the E1443D, E1443Q and R881D/D1432R showed evidence of reduced stability relative to the wild-type model, whilst the W1442Y mutant was most similar to the wild-type channel (Figure 7A and B). This is consistent with the electrophysiological data for the human W1440Y mutant, which shows no significant differences with the wild-type channel in any gating parameter, including peak current.

**Figure 7.**
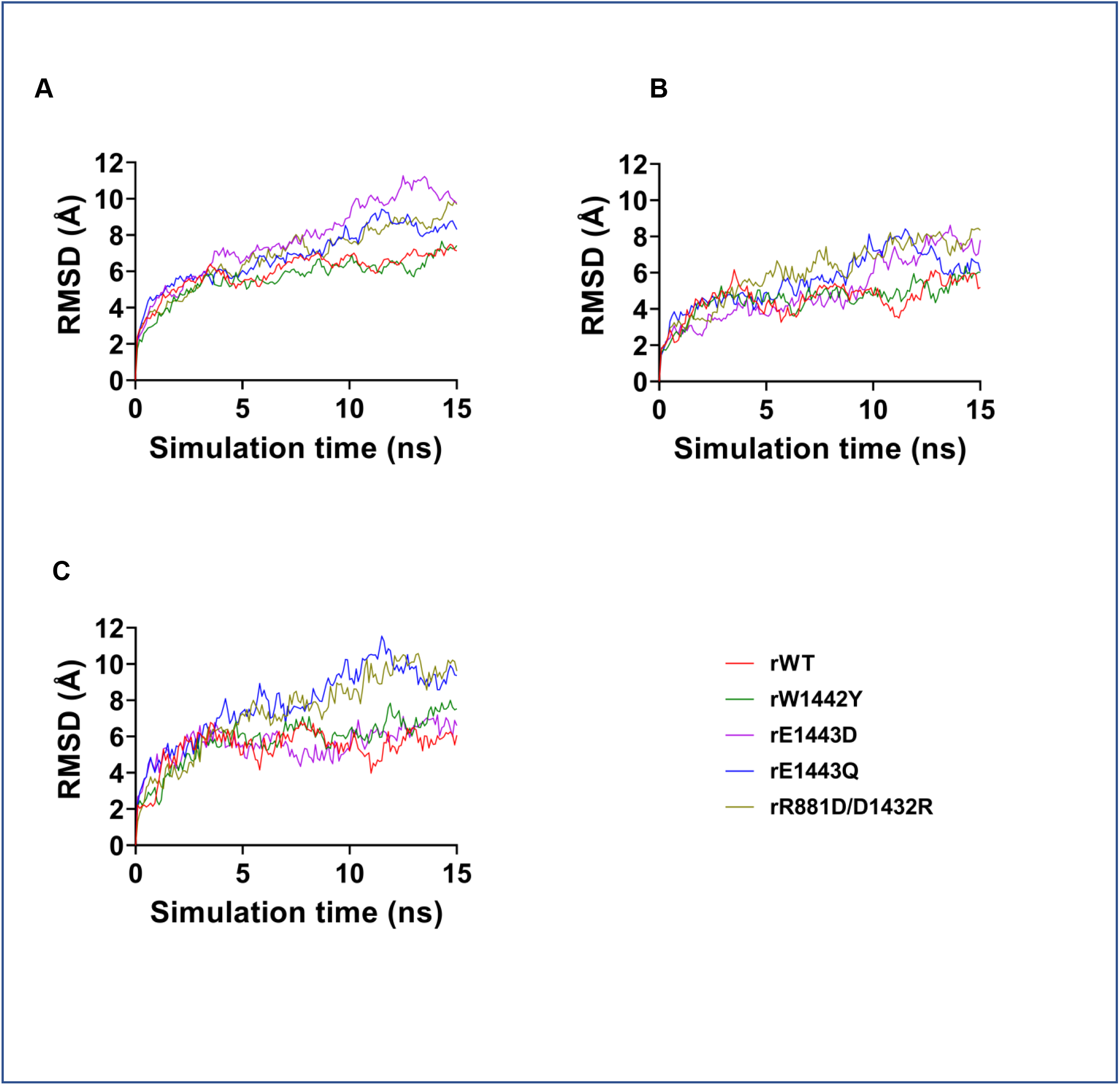
MD simulations of the global and local stabilities of the simulated rat Nav1.5 channels. RMSD values of C(α) atoms of the wild-type (WT) and mutant rat (r) Nav1.5 channels. The first frame of the MD simulation as a function of time was examined for, **(A)** all residues in the protein, **(B)** residues of the S5-S6 helices lining the pore of the channel and **(C)** residues of the complex salt-bridge and cation-π bonds connecting the DII-DIII turret interface.

The RMSD backbone values for the turret interface residues (Figure 7C) provided a finer-grained analysis. For example, the flexibility of the E1443D mutant was comparable to the wild-type and the W1442Y mutant and was now distinct from the higher flexibility shown by the R881D/D1432R and E1443Q mutations. The human versions of R881D/D1432R and E1443Q were devoid of any detectable gating behavior, whilst the human version of E1443D retained some peak current activity (Figure 2A, Table 2). Hence, the modeled stability of the turret interface region provides the clearest relationship with function, even in the presence of some global destabilization.

### Mutant-specific destabilization of the turret interface

To better understand how the mutations affect the local stability at the DII-DIII turret interface, we applied MD simulations to calculate both energy and distance values for the rat R881-E1443 and R881-D1432 salt-bridges and their cation-π interactions with W1442 and Y1428. Firstly, we analysed these values across the entire time course of the simulation, to trace their evolution (Figure 8A). Secondly, for each bond simulated, we plotted the energy against distance values from the last 5 ns of the simulation. The resulting cluster diagrams emphasize, for each of the mutants, both the local dispersion of these values and their deviation from the wild-type values (Figure 8B).

**Figure 8.**
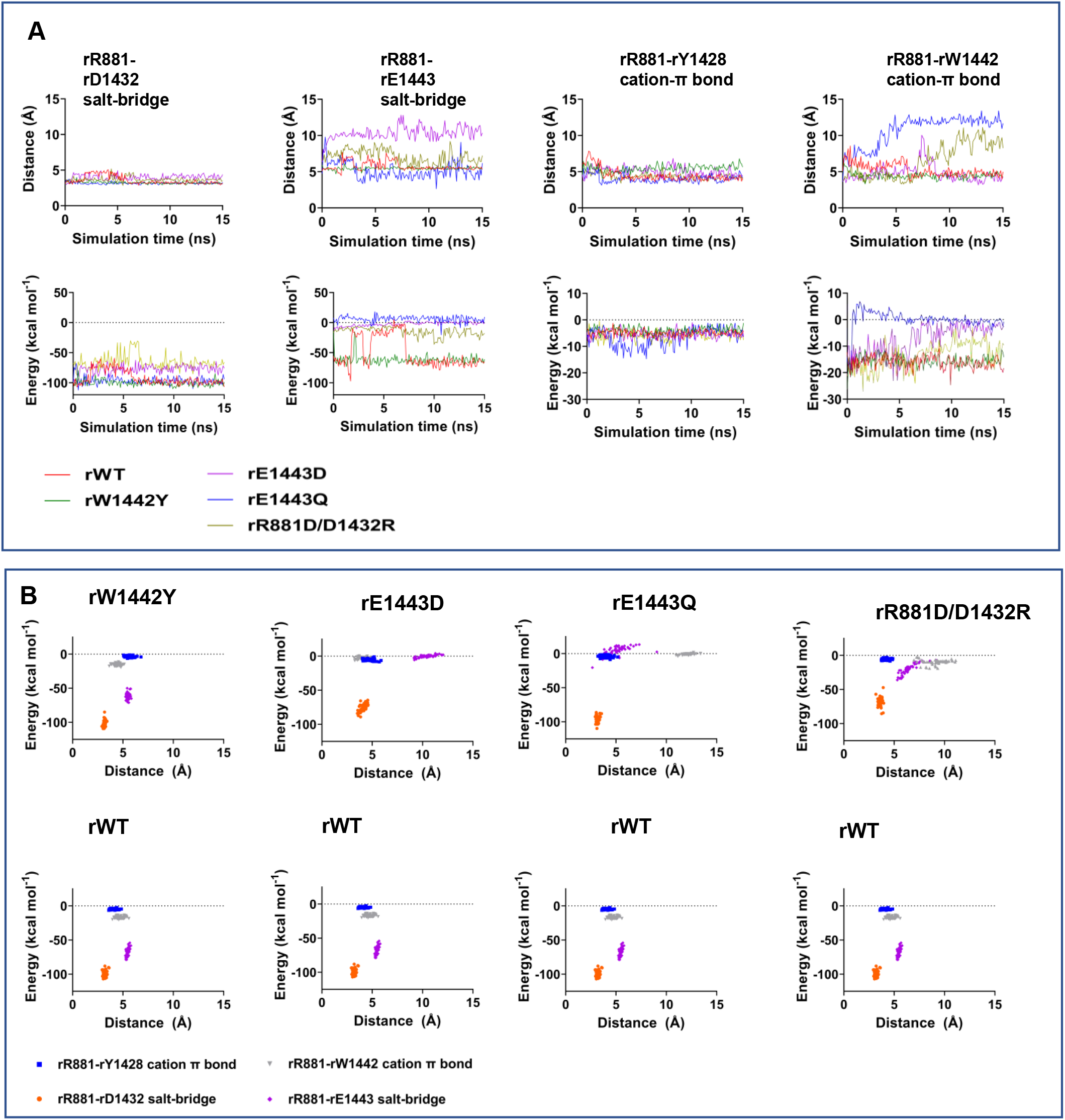
Comparative analyses of energy values and distances for specific pairs of residues in the DII-DIII turret interfaces of the simulated rat Nav1.5 channels. **(A)** Graphical depiction of the interactions studied across the entire time course of the simulation for the wild-type (WT) and mutant rat (r) Nav1.5 channels. Energy and distance values are presented for every 0.1 ns frame of the MD simulation as indicated. **(B)** Comparative cluster plots of energy values compared to distances, between pairs of residues in the DII-DIII turret interface, for each mutant, individually compared to the wild-type channel as indicated. Note: for presentational clarity, the numbers refer to the wild-type structure, although the identity of the residues may differ for the individual mutants.

The bond energies and distances exhibited by the W1442Y mutant were almost indistinguishable from the wild-type (Figure 8A and B). This is again consistent with the electrophysiological data showing no differences in gating behavior for the human W1440Y mutant compared to the human wild-type Nav1.5 (Table 2). For the case of the modeled E1443D mutant, the salt bridge it formed with R881 was weakened compared to the wild-type protein. This occurred despite E1443D retaining a negatively charged residue. The E1443D mutant also slightly reduced the stability of the salt-bridge between residues R881 and D1432 and the cation-π interaction between R881 and W1442. However, the rat E1443Q mutant (equivalent to the human BrS mutant E1441Q), severely perturbed the interaction between residue 1443 an R881 and now also disrupted the R881-W1442 cation-π bond, to the point where it was unlikely to be maintained. The salt-swapped mutant, (R881D/D1432R), exhibited both higher energy and increased distance profiles for the ionic interactions within the complex salt-bridge and also a perturbed the R881-W1442 cation-π bond. These results, when correlated with the electrophysiological data for the equivalent human mutations, suggest that the salt-bridge between E1443 and R881 and the cation-π bond between R881 and W1442 are the most important in maintaining a functionally permissive structure to the DII-DIII extracellular loop regions. In contrast, the R881-Y1428 cation-π bond was relatively unaffected by any of the modeled mutations (Figure 8A and B).

### Mutations in the turret interface perturb the geometry of the DEKA selectivity ring, reduce the radius of the pore and reduce sodium ion penetration

We hypothesize that the mutation-induced instability within the DII-DIII turret region may allosterically distort the selectivity filter and/or the inner pore geometry, thereby inhibiting sodium ion flux. We calculated the distance between each pair of residues in the selectivity ring of the wild-type and mutant channels using MD simulation. Here, the W1442Y mutant exhibits a DEKA ring geometry only slightly different from that in the wild-type Nav1.5. By contrast, each of the other mutations displays distinct and shorter distance geometries compared to the wild-type channel (Figure 9A and B). The effects of these changes on sodium ion penetration were studied using simulations in 150 mM NaCl. The positions of individual sodium ions within the outer vestibule of the pore were tracked for the wild-type and each of the mutants over 15 ns (Figure 9C; Supplemental Figures 1 A-E). In all cases, sodium ion diffusion along the z-axis became constrained within about 2 ns. In the case of the wild-type simulation, the sodium ion co-localized with the position of the DEKA ring (Figure 9C). A closer examination of the simulation revealed that a sodium ion was transiently chelated by the two DEKA acidic residues D373 and E901, before moving into the inner vestibule, to be replaced by a second incoming sodium ion (Figure 9D; Supplemental Figure 1A). This is broadly consistent with previous MD simulations and evidence from the cryo-EM structure of the cockroach Nav channel (28, 30, 31). Interestingly, however, in the mutants, the sodium ion predominantly remained in the outer vestibule, constrained between 10 and 15 Å above the DEKA ring (Figure 9C). This suggests that in the mutants, the diffusion of the sodium ions through the outer vestibule was compromised compared to the wild-type channel (Figure 7C).

**Figure 9.**
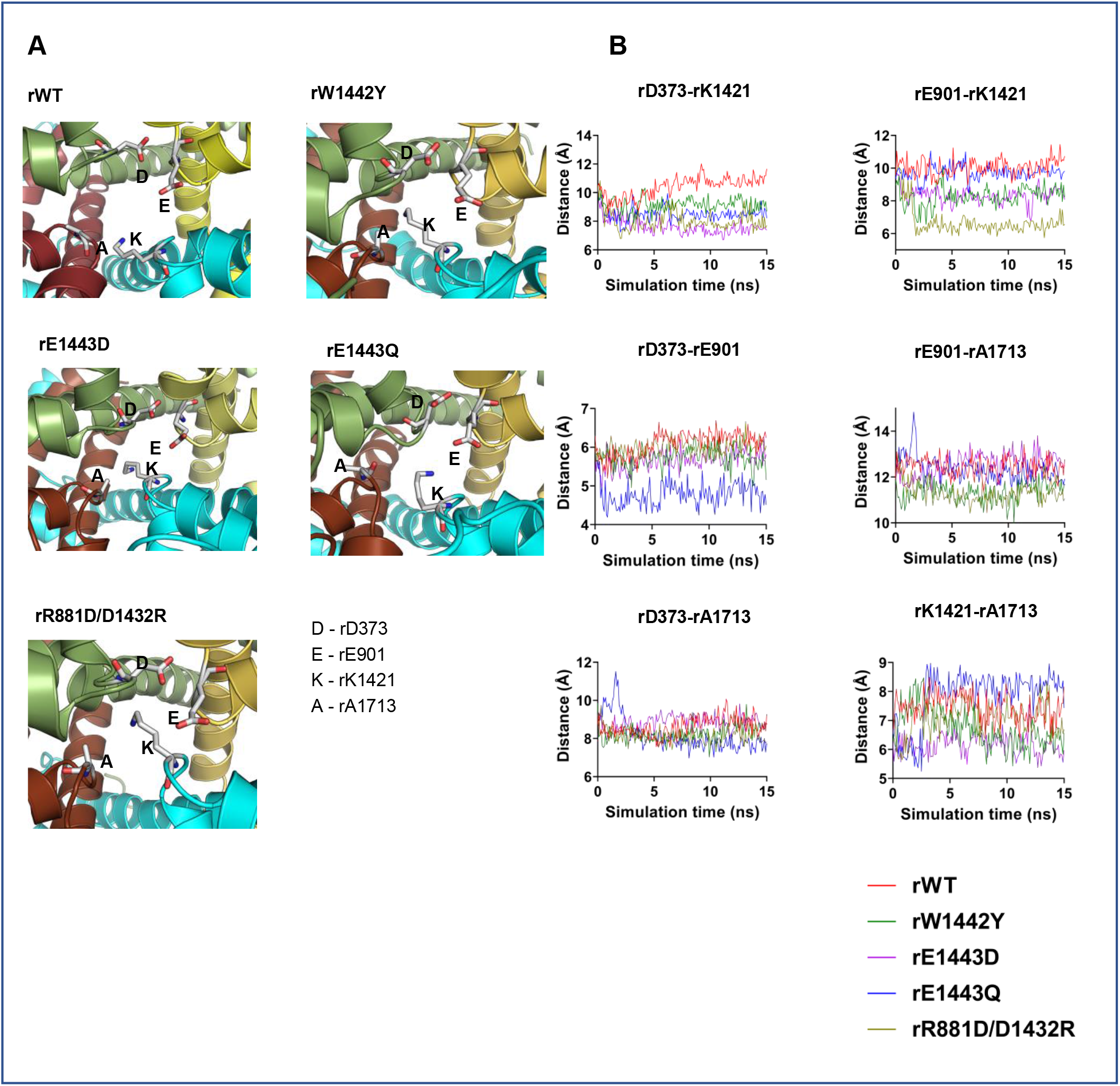
Effect of the DII-DIII turret mutations on the geometry of the DEKA selectivity ring of the simulated rat Nav1.5 channels. **(A)** Diagrammatic representation of the contraction of the DEKA selectivity ring at 15 ns, for the simulated wild-type (WT) and mutant rat (r) Nav1.5 channels, as indicated. **(B)** Distances between pairs of residues in the DEKA ring, as indicated, equilibrated throughout a 15 ns simulation.

**Figure 9.**
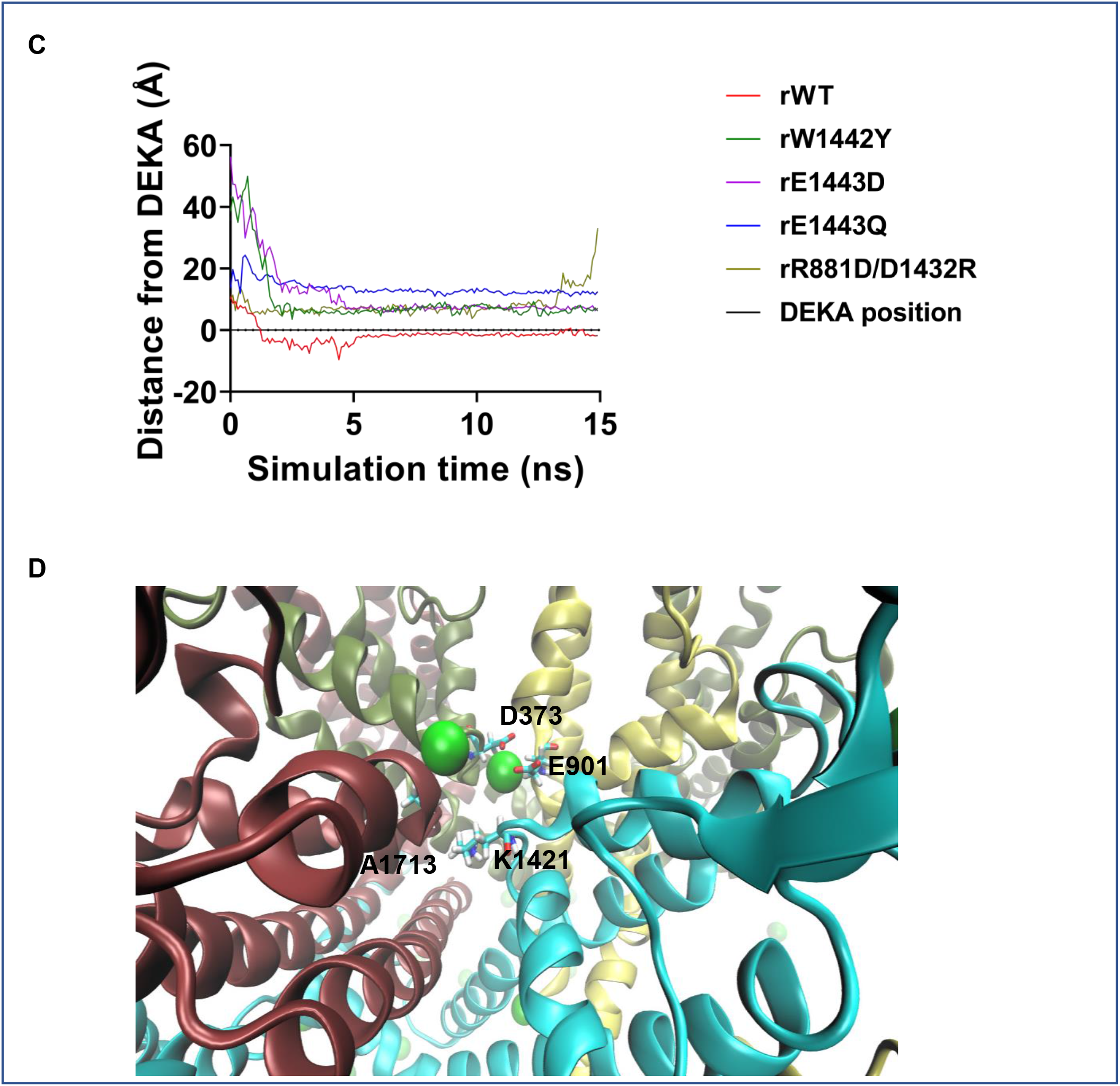
Effect of the DII-DIII turret mutations on the geometry of the DEKA selectivity ring of the simulated rat Nav1.5 channels. **(C)** Tracking the position of sodium ions in the outer vestibule of the channel before they travel down the pore. The dotted line corresponds to the mean position of the DEKA residues along the z-axis and the traces represent the position of the sodium ions along the z-axis across the time course of the simulation. **(D)** Frame from the wild-type simulation (**Supplemental Figure 1**), showing a sodium ion transiently trapped between the D373 and E901 residues of the DEKA selectivity ring, with their negatively charged carboxylate groups facing the ion.

We next modeled the pore topology, for each structure after 15 ns simulation, from the DEKA ring to the intracellular activation gate, using two computational approaches. Firstly, we used the MOLE 2 program (32) to provide a pseudo-three dimensional rendering of the pore profile (Figure 10A). Secondly, we calculated the pore radius along its z-axis (Figure 10B). In the wild-type structure, the DEKA constriction point and the inner vestibule (4) were clearly identified (Figure 10A and B). In contrast, the E1443D and E1443Q mutants both showed a reduced pore radius throughout the length of the channel, but with the E1443Q mutant showing a higher degree of constriction than the E1443D mutant (Figure 10B). This supports the electrophysiological results that the human equivalent mutant E1441D, shows a reduced but still detectable conductance, whilst the human E1441Q BrS mutant has no detectable electrophysiological activity (Figure 2A). Strikingly, we could not compute any pore through the modeled rat R881D/D1432R mutant, using either approach (Figure 10A and B). Hence, the occlusion of sodium is predicted to be particularly severe in this mutant and is consistent with the complete absence of gating activity for the human equivalent mutant.

**Figure 10.**
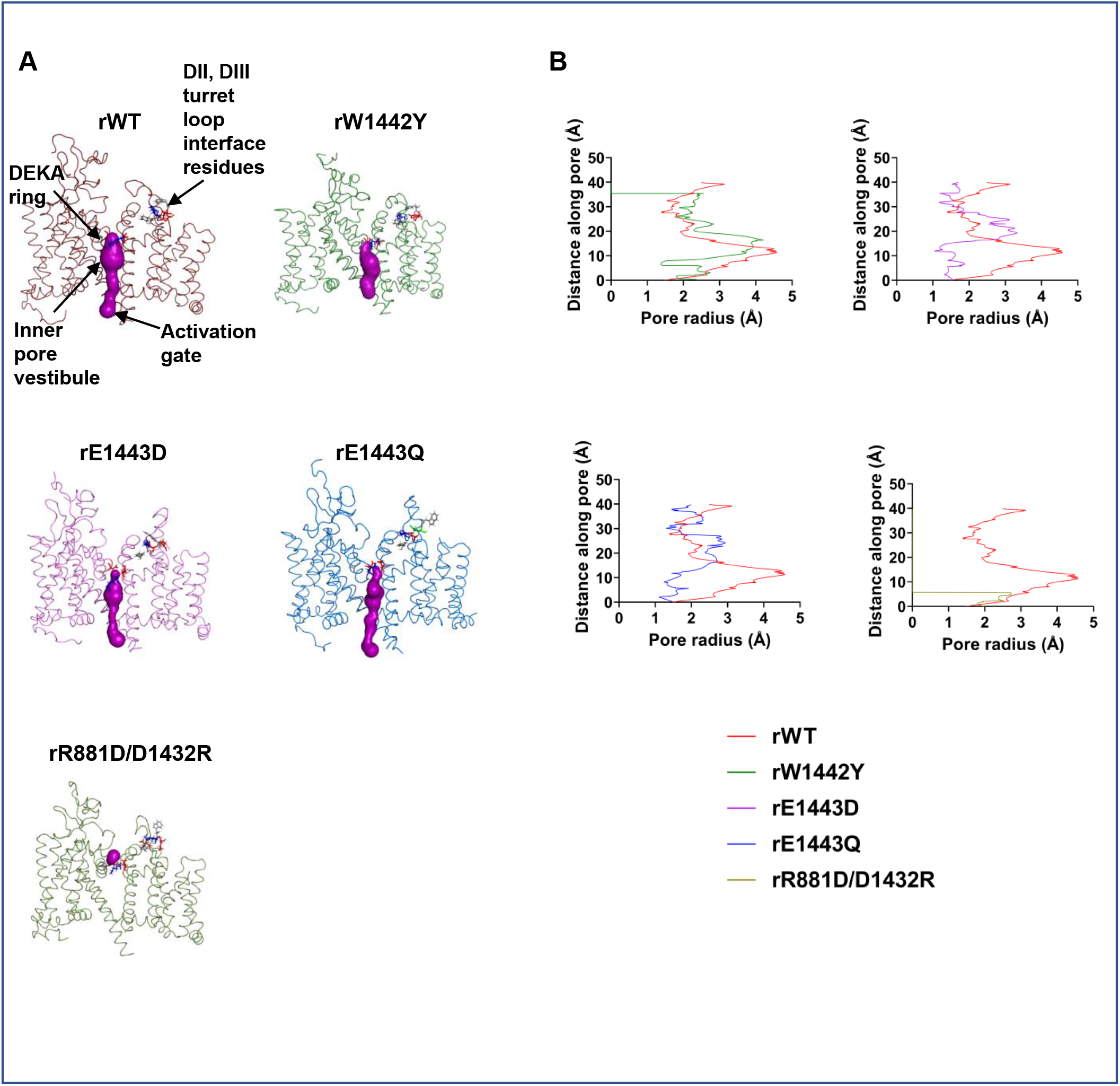
Effect of the DII-DIII turret mutations on pore geometry of the simulated rat Nav1.5 channels. **(A)** Cross-section of the last-frame of each MD simulation for wild type (WT) and mutant rat (r) Nav1.5 channels as indicated. Pore profiles, as predicted using MOLE 2 are highlighted in purple. The locations of key regions within the pore are indicated for the wild-type channel. The DEKA ring and DII-DIII turret interface residues are highlighted as stick format for wild-type and each mutant. **(B)** Pore radius profiles for the simulated wild-type (WT) and mutant rat (r) Nav1.5 channels are as indicated and computed to the intracellular surface of the channel. The distance along the pore starts from the DEKA selectivity ring (0) to the intracellular activation gate (50Å).

## Discussion

In this work, we highlight a conserved, complex salt-bridge with associated cation-π bonds, that connects the DII and DIII extracellular turret loops and helps encase the Nav1.5 outer vestibule. Using the combined approaches of mutagenesis, electrophysiology and MD simulations, we show that targeted mutations to these residues significantly compromise Nav1.5 gating behavior and provide new structural insights into the pathological human BrS/SND mutations that occur in this region (11, 13).

All the human mutants that we have studied, including the mutants that displayed no activity in electrophysiological recordings, were nevertheless expressed on the plasma membrane to the same extent as the wild-type channel (Figure 5). The previously reported human D1430N BrS-associated null-mutant is similarly expressed on the plasma membrane (12). There are multiple and extensive quality-control mechanisms within the secretory pathway that can efficiently detect and destroy aberrantly-folded proteins before they reach the plasma membrane (33). Hence, the compromised gating behavior of the mutant Nav1.5 channels is unlikely to be explained by a large-scale misfolding of the channels. Rather, the mutations are more likely to introduce relatively subtle structural changes, which block one or a small number of gating steps.

For the human mutants that retained some gating activity, only the peak currents and steady-state activation parameters were significantly different from those of the wild-type channel. Steady-state inactivation parameters and recovery from inactivation kinetics were not affected (Table 2; Table 3). Taken together with the failure of veratridine to rescue the null-mutants, this suggests that disruption of the extracellular DII-DIII interface on the outer vestibule turrets selectively inhibits one or more of the conformational changes that underlie channel activation. Yet channel activation depends on conformational changes that occur deeper within the pore and within the peripheral voltage-sensors, which are physically distant from the DII-DIII interface on the extracellular turret (2). Hence, the DII-DIII mutations are likely to induce longer-range, allosteric perturbations.

Complex salt-bridges play an important role in stabilizing secondary structures and adjacent domains within proteins (34). The appropriate ionic contacts between residues of the complex salt-bridge depend not only on their charge but also on their precise geometry and orientation. In these situations, arginine is significantly more common than lysine as the cation partner. This is because the delocalized positive charge on the arginine guanidinium group can simultaneously facilitate multiple electrostatic interactions (14). Thus, in the wild-type rat Nav1.5 structure, the η1 and η2 nitrogens of the R881 guanidinium group individually contact the carboxylate groups of residues D1432 and E1443 respectively, whilst the ∊ nitrogen lies over the aromatic group of Y1428, contributing to the cation-π interaction (Figure 2B). The positive charge on a lysine side chain cannot do this as easily because it is more tightly localised. These considerations probably explain why even lysine cannot functionally substitute for residue R878 in human Nav1.5 (11).

Of the nine null-mutations studied, seven were targeted to the complex salt-bridge. Of the six mutants that retained some gating activity, five were targeted to the cation-π bonds (Figure 2A). Thus, the complex salt-bridge is likely to provide the major quantitative stabilization for the DII-DIII turret interface, whilst the cation-π bonds contribute secondary, although still functionally important reinforcement. The MD simulations enable a further examination of these interactions. For example, in the modeled rat E1443Q mutant (equivalent to the putative human BrS null-mutant E1441Q, Table 1), the normal salt-bridge between residues 881 and 1443 cannot form. This effectively abolished the cation-π bond between residues R881 and W1442 and yet the salt-bridge between residues R881 and D1432 and the cation-π bond between R881 and Y1428 were unaffected (Figure 8A and B). The second null-mutant examined by MD simulation was R881D/D1432R, in which the charged residues R881 and D1432 were reciprocally swapped. This arrangement weakened but did not eliminate, the normal salt-bridge between these two residues. Strikingly, however, the R881-E1443 salt-bridge as well as the R881-W1442 cation-π bond were largely abolished (Figure 8A and B). A close examination of the final equilibrium residue positions from the relevant MD simulation for the R881D/D1432R mutant suggests a possible explanation. Here the new position of the guanidinium group on the repositioned R1432 residue can maintain a hydrogen bond to the D881 residue but is now too far away from the E1443 residue to make a strong contact. In consequence, both the E1443 and the W1442 residues can no longer be constrained (Supplemental Figure 2A).

The individual replacement of each of the aromatic residues, human W1440 and Y1426 with alanine significantly reduced peak current, but with no significant effects on steady-state gating or kinetic parameters. However, the replacement of human W1440 with tyrosine showed no significant gating effects compared to wild-type human Nav1.5 (Table 2). Neither were there any major differences in the structural stabilities of the modeled rat wild-type Nav1.5 and W1442Y mutant (Figure 8; Figure 9; Supplemental Figure 1). Thus, the precise form of the aromatic amino acid at this position is not critical and indeed, tyrosine not tryptophan is present at this position in most Nav channel isoforms (Supplemental Figure 2C). By contrast, there are more constraints on the second cation-π-forming aromatic residue (human Y1426/rat Y1428). For example, even the replacement of this tyrosine with phenylalanine reduced peak current by about 50% for the human mutant (Table 2). This suggests that the tyrosine hydroxyl group is functionally important and in the rat structure, Y1428 forms a hydrogen bond with the DII W882 residue (Figure 2B), thus holding either or both the W882 and Y1428 residues in a constrained orientation that may further stabilize the DII DIII turret interface.

To investigate the effect of the turret mutations on the DEKA ring and pore geometry, we performed additional MD simulations. In the channel structure on which these simulations were based, the cytoplasmic inactivation gate (Figure 1A) is engaged with its DIV binding site and the intracellular pore activation gate is closed. The structure is therefore thought to represent the inactivated state (2), and so we do not expect the sodium ions to fully traverse the channel in this conformation. However, given the differences in pore geometries between the wild-type and mutant structures, important insights are still likely to be obtained. For example, previous MD simulations of Nav channels have typically used inactivated state structures as templates (35).

The DEKA ring acts as a constriction point and selectively favors sodium ion permeability over other cations. Site-directed mutagenesis experiments have identified an important role both for the acidic groups and the lysine residue of the DEKA ring (36, 37). The cryo-EM structure of the cockroach Nav channel, solved in the presence of 300 mM NaCl contains a region of electron density consistent with a largely dehydrated sodium ion, coordinated by the two DEKA acidic residues (38). Previous MD simulations of the mammalian Nav channel DEKA rings used a homology modeling approach but used prokaryotic sodium channels as templates. These simulations suggest that the DIII lysine residue acts in part to repel an incoming sodium ion and facilitate its transient capture by the adjacent DI and DII acidic residues, whose relative positions favor a sodium ion compared to a larger potassium ion (28, 30). It is proposed that a second, incoming sodium ion displaces the captured sodium ion, which is then free to pass through the DEKA constriction site (28). In these previous MD simulations, the protonated lysine residue adopted two distinct conformational categories: an ‘off-state’, in which the lysine side chain interacts with the adjacent DII glutamate residue and blocks the pore; and an ‘on-state’, in which the lysine side chain is sequestered perpendicularly, to allow sodium ion permeation (28, 30). Our MD simulations were based on the rat Nav1.5 structure. They predict that in the wild-type Nav1.5, the K1421 residue predominant adopts a perpendicular, ‘on’ conformation, allowing unobstructed access of the sodium ions for the pore (Figure 9A and B). In the permeation simulations of the mutants, the misalignments of the K1421, D373 and E901 residues and possible steric interference from the enhanced flexibility of the DII-DIII turret, could explain why sodium ions were largely restricted to locations above the DEKA ring (Figure 9C; Supplemental Figure 1 A-E). A particularly striking example was the rat R881D/D1432R mutant. Here, the K1421 residue predominantly adopted a less perpendicular conformation, reminiscent of the ‘off’ states (28, 30), such that the lysine side chain lies across the pore partially obstructing it (Figure 9A). Of all the simulated mutants examined, the rat W1442Y showed the least discrepancy in DEKA ring and pore geometry compared to the wild-type channel and the electrophysiological properties of the human equivalent mutant were not significantly different from the human wild-type channel (Table 2). Nevertheless, in the simulation, the movement of sodium ions was largely constrained to the outer vestibule and above the DEKA ring (Figure 9C). A clearer understanding of this observation may require a further comparison between open and inactive channel state conformations.

It is conceivable that perturbation of the DEKA ring may relax ion selectivity. However, this is unlikely to be the case for those human mutants that display a complete absence of current gating since the external bath solution contained 2 mM KCl and 1.5 mM CaCl_2_. For those human Nav1.5 channels that retained some gating activity, the ion-selectivity of the channel can be investigated by comparing their reversal potentials (36, 39). Importantly, the reversal potentials of the functional mutants were not significantly different from the wild-type value (Figure 2A). This supports the view that the DEKA rings in each of the active human mutants retained selectivity for sodium ions, despite their altered geometry.

The most dramatic changes predicted from the MD simulations occurred in the dimensions, geometries and stabilities of the inner pore, comprising the S5 and S6 helices from each domain. In particular, the increased pore radius corresponding to the inner vestibule was prominent in the rat wild-type and W1442Y mutants. In contrast, the inner vestibule was significantly attenuated in the rat E1443Q and absent altogether in the rat R881D/D1432R mutant (Figure 10A and B). Both equivalent human mutations were electrophysiologically inactive (Figure 2A). This attenuation in the inner vestibule radius was less dramatic in the modeled rat E1443D mutant (whose human equivalent retained about 6% of the wild-type peak current activity) but was still noticeable compared to the wild-type and W1442Y profiles (Figure 10A and B). Hence, the degree of pore constriction in the MD simulations for the mutants – particularly within the inner vestibule – broadly correlated with the gating activity of the equivalent human mutants.

Except for the human W1442Y and Y1426F mutants, all other human activity-retaining mutants exhibited significantly higher values of the slope-factor, *k* (activation) compared to the wild-type channel (Table 2). This slope factor is inversely proportional to the effective charge (*z*) transferred across the membrane during activation (40, 41). The larger values of *k* (activation) therefore imply that the voltage-sensing movements in the VSDs of the mutants are restricted relative to the wild-type channel. These movements are normally tightly coupled to pore opening (4). Each of the four S6 helices lining the inner pore vestibule contains a highly conserved asparagine residue, that is essential for this coupling. However, the underlying molecular mechanism is unclear (42). In rat Nav1.5, these residues are: N407 (DI); N935 (DII); N1465 (DIII) and N1767 (DIV). We note that in the rat MD simulations, the displacements between the asparagine residues of the wild-type and W1442Y structures lie between 1.99 and 3.55Å. In contrast, the asparagine residues on the S6 helices from the rat E1443D, E1443Q and R881D/D1432R mutants show a much greater displacement of up to 8.71Å, relative to the wild-type channel (Supplemental Figure 2B). It is possible that in these mutants, the misalignment of the critical asparagine residues obstructs the opening of the pore, which in turn, restricts the movement of the VSD S4 helices. A rigorous test of this hypothesis will require significant further work. For example, the application of biophysical methods that can monitor the S4 helix movements within the individual VSDs of each mutant (21, 43).

The presence of a similar cation-π bonded complex salt-bridge at the DII-DIII turret interface in Nav1.2, Nav1.4 and Nav1.7 (31, 44, 45), together with their strong sequence conservation in all Nav channel isoforms (Supplemental Figure 3A), suggests that all Nav channel isoforms will display this structural feature.

Interestingly, several independent missense mutations corresponding to these residues in human Nav1.1 are associated with inherited epileptic encephalopathy (46) (Supplemental Figure 3A). In addition, an arginine to glutamine missense null-mutation in human Nav1.7 is associated with congenital insensitivity to pain. This Nav1.7 residue is designated R896 (47) or R907 (46), depending on the alternatively spliced Nav1.7 isoform, and is equivalent to human Nav1.5 R878 (Supplemental Figure 3 A and B). Finally, eukaryotic voltage-gated sodium channels and voltage-gated calcium channels are derived from a common ancestor by gene duplication (48). The Cav1.1 family of voltage-gated calcium channels similarly contains a salt-bridge, bounded by likely cation-π bonds at the DII and DIII extracellular turret interface (Supplemental Figure 3C). To our knowledge, there are no clinical reports of mutations affecting these Cav1.1 residues. However, if such mutations are discovered, we predict that the channels will either be non-functional or severely compromised and are likely to have pathological consequences. Hence our current work provides general insights, not just into Nav1.5-associated cardiopathologies but also into a variety of distinct and previously unconnected ion-channelopathies.

## Methods

### DNA constructs and mutagenesis

Bicistronic plasmids encoding hNa_v_1.5 (pIRES2-EGFP-hNav1.5; also called hH1, accession no. M77235) and the R878C mutant (pIRES2-EGFP-R878C) together with a separately expressed EGFP were a gift from Dr Ming Lei (University of Oxford, UK). All further mutations were introduced by site-directed mutagenesis (QuikChange Lightning kit, Agilent) according to the manufacturer’s instructions and checked by DNA sequencing.

### Cell culture and transient transfections

Human embryonic kidney (HEK293F) cells were cultured in DMEM/F-12 GlutaMAX medium (Invitrogen) supplemented with 10% fetal bovine serum (Sigma–Aldrich) at 37 °C and 5% CO_2_. For patch-clamp experiments, 0.5 μg of Na_v_1.5 wild-type (WT) or appropriate mutant were transiently transfected using polyethylenimine (PEI, 1 μg/μl) at a DNA/PEI ratio of 1:4, on 18 mm coverslips in 6-well plates. For the biotinylation assays, HEK293F cells were transiently transfected in 100 mm dishes with 8 μg of Na_v_1.5 WT, mutant or empty vector pEGFP-N1 as a negative control, using a DNA/PEI ratio of 1:3.

### Cell surface biotinylation assay

Biotinylation assay was adapted from (22) with slight modifications. 48h after transiently transfecting HEK293F cells with appropriate plasmids, cells were washed twice with cold 1X PBS solution, then incubated with freshly prepared 0.5 mg/ml EZ-link™ Sulfo-NHS-SS-Biotin (Thermo Scientific) in cold 1X PBS for 15 min at 4 °C. To quench and remove excess biotin the cells were washed twice with 200 mM glycine (in cold 1X PBS), and then twice with cold 1X PBS. Cells were then lysed in 1 ml of lysis buffer (50 mM HEPES pH 7.4; 150 mM NaCl; 1.5 mM MgCl_2_; 1 mM EGTA pH 8; 10% Glycerol; 1% Triton X-100) supplemented with 1X Complete Protease Inhibitor Cocktail (Roche, Sigma-Alrdrich) at 4°C for 1 h, and then clarified by centrifugation; 16,000 X g for 15 min at 4 °C. 500 μg of the supernatant was incubated with 50 μl Pierce™ Streptavidin Agarose beads (Thermo Scientific) overnight at 4 °C, while the remaining supernatant volume was kept as input fraction. Following the overnight incubation, the Streptavidin beads were pelleted at 1000 X g for 1 min and washed thrice with cold 1X lysis buffer and then eluted with 50 μl of NuPAGE LDS Sample Buffer (4X; Invitrogen) supplemented with 100 mM DTT at 37 °C for 30 min. The input fractions were similarly treated in LDS buffer + 100 mM DTT at equal final concentrations to function as a loading control indicating the total Nav1.5 expression, while the biotinylated fractions represent just the Na_v_1.5 expression at the cell surface.

### Western blotting

The input (total Nav1.5 expression) and biotinylated (cell surface-expressed Nav1.5) samples were separated on NuPAGE 4–12% Bis-Tris gels (Invitrogen, ThermoFisher) and transferred to nitrocellulose membrane (iBlot transfer system, Invitrogen, ThermoFisher). Membranes were blocked with 5% milk in TBS-Tween (0.1%) for 45 mins at room temperature before overnight incubation of primary antibody at 4 °C. Primary rabbit polyclonal anti-(human Na_v_1.5), (residues: 1978-2016) (#ASC-013, Alomone Labs) and rabbit polyclonal anti-(actin) (A2066, Sigma), both used at a concentration of 1:200 were followed by a one-hour incubation with secondary Goat anti-(Rabbit IgG-HRP conjugate) (#1706515, Bio-Rad) at a concentration of 1:8000 at room temperature. Amersham ECL Western Blotting Detection Reagent (GE Healthcare) was used to activate the HRP and the chemiluminescent signal detected on Amersham Hyperfilm ECL (GE Healthcare). Immunoblot intensities were quantitated using Image J and normalized against their loading controls.

### Electrophysiology

Electrophysiological experiments were carried out as previously described (21) with slight modifications. GFP fluorescence from pIRES vector helped visualize the efficiently transfected cells chosen for the whole-cell patch-clamp recordings which were made 48h post-transfection. For all measurements, of inactivation, activation and recovery from inactivation, the whole-cell configuration of the patch-clamping technique was employed using an Axopatch 200 amplifier (Axon Instruments, USA), Digidata 1322A digitizer (Axon Instruments, USA), and the Strathclyde Electrophysiology Software Package (WinWCP, Department of Physiology and Pharmacology, University of Strathclyde). All the experiments were performed at room temperature (20-22°C). The extracellular bath solution used to superfuse the cells was set at a flow-rate of 2–3 ml min^−1^ and was composed of 60 mM NaCl, 2 mM KCl, 1.5 mM CaCl_2_, 10 mM glucose, 1 mM MgCl_2_, 90 mM CsCl_2_, 10 mM HEPES, pH 7.40 ± 0.01 adjusted with NaOH. The internal solution contained 35 mM NaCl, 105 mM CsF, 10mM EGTA, 10 mM HEPES, pH 7.40 ± 0.01 adjusted with CsOH and filter sterilized (0.22 μm). Patch pipettes were prepared from borosilicate glass capillaries (1.2mm OD, Harvard Apparatus) using electrode puller model P87 (Sutter Instrument Co., USA) with pipette resistance maintained between 1.5-2.5 MΩ. Signals were low-pass Bessel filtered at a cut-off frequency of 5 kHz and sampled at 125 kHz. Series resistance compensation (70–85%) was set for each recording, and leak currents were subtracted using a P/4 protocol. All recordings were made after allowing at least 3 minutes after membrane rupture. The liquid junction potential (typically close to 2 mV) was not corrected for and data from cells with a clear loss of voltage control as demonstrated by poor I/V relationships were removed.

### Voltage protocols

All protocols employed a holding potential of - 120 mV. The voltage dependences of steady-state activation and inactivation were examined using a 100 ms duration pulse, to activating test voltages ranging between - 140 mV to + 45 mV in 5 mV increments with each sweep, followed immediately by a pulse to a fixed voltage of −40 mV, of 50 ms duration. For measurements of steady-state inactivation, peak currents elicited by each of the latter −40 mV pulses, were normalized to the maximum peak current generated and then plotted against the voltage of the preceding 100 ms conditioning pulse. The data assumed a sigmoid function, decreasing with positive shifts of membrane potential. For the activation protocol, peak current densities (pA/pF) were calculated by dividing the peak current amplitude by whole-cell capacitance and the current-voltage relationship plotted. Both inactivation and activation data were fitted with a single Boltzmann function:

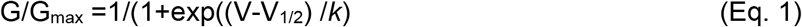

where G/G_max_ is the normalized conductance or current, V_1⁄2_ is the voltage of half-maximal activation or inactivation, *k* is the slope factor, and V is the test voltage or conditioning voltage. The G terms for each voltage V, were calculated from observed peak currents I_Na_ and their reversal potential E_Na_ observed at large depolarizing activating test voltages as:

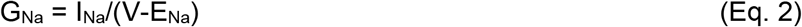

The recovery from inactivation protocol employed a standard two-pulse protocol (P1) and (P2) both at - 40 mV for 50 ms but separated by a return to the holding potential (−120mV) of varying time courses between 1 to 200 ms. The ratio P2/P1 was then calculated to normalize the peak current, which indicated the fraction of current available for activation after depolarizations at varying durations. These plots were fitted with a double exponential function as follows:

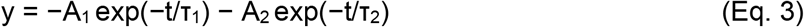

Where A_1_ and A_2_ are the amplitudes of the fast and slow components respectively, τ_1_ and τ_2_ are the time constants of recovery from inactivation for the fast and slow components respectively and t is time.

### Application of veratridine

100 μM veratridine (Abcam, ab120279) was applied in the bath solution and continuously perfused during the whole-cell recording of HEK293F cells transiently transfected with the wild-type Nav1.5 or those mutants which showed no detectable activation current (49). Veratridine was allowed at least 5 minutes to perfuse the bath before starting the recording. Effects of washing out the drug were not performed.

### Disulfide locking experiment

Double cysteine mutations were carried out on the charged residues likely taking part in salt bridge formation (human R878C/E1441C and R878C/D1430C). Whole-cell recordings of HEK293F cells transfected with the wild-type Nav1.5 or either of these two mutants were performed after continuously perfusing the bath solution with freshly prepared 1 mM β-mercaptoethanol (βME), allowing at least 5 minutes before the start of recording (20).

### Molecular dynamics simulations

The cryo-EM structure for rat Nav1.5 (PDB ID: 6UZ3) (2) was used for molecular dynamics simulations. A few residues were not well-resolved in the structure and so were not modeled. These include residues: 213, 291, 298-303, 799-805 as well as residues: 425-698 and 945-1189, which correspond to intracellular domains of the channel. MD simulations were performed using NAMD 2.9 (50) with CHARMM36 force fields (51). All simulations were conducted at 310 K and 1 atm. For the wild-type structure, simulations were run on the original PDB structure: 6UZ3. Each of the mutants was engineered using the “Mutator” plugin of VMD 1.9.4 (29).

To model the dynamics of each system, the protein was embedded into a lipid bilayer. Each protein was inserted into the palmitoyl-oleoyl-phosphatidylcholine (POPC) bilayer (containing 229 lipid molecules), which was then solvated by TIP3P waters (~ 36000 water molecules) and neutralized by 150 mM NaCl (50, 51). Simulations were preceded by pre-equilibration steps. Firstly, the system was energy-minimized for 2000 steps and then equilibrated using a time step of 1 fs for 0.5 ns. To relax the lipid-protein interactions, all atoms were constrained except the lipid tails. Non-bonded Fix (NBFIX) parameters were utilised for this step as they are optimized for the specific case of ions interacting with carbonyl oxygen atoms (52). The next step of pre-equilibration was a minimization run, that lets the system move to the lowest energy conformation. This was followed by equilibration with the constrained protein so the environment can relax. This was also energy minimized for 0.2 ns run and equilibrated for 0.5 ns at a time step of 1 fs. These steps ensured the simulation ran smoothly and the starting velocity of atoms was not too high. Subsequently, productive simulations were run for a duration of 15 ns with the same time step of 1 fs.

### Analysis

On successfully running the simulations, trajectory files (with a dcd extension) were generated and subsequently analysed to investigate properties of the channel and region of interest. The RMSD values were generated through VMD’s “RMSD Visualiser” tool which generated a value for every fs. VMD was used in the analysis of the DEKA ring. With VMD’s ‘Tkconsole’ (53), a tcl script was run to compute the atomic distances between the alpha carbons of the residues for every 1000 frames. For the distances between residues of interest, distances between relevant charged groups on each residue was used. Correspondingly, to calculate the energy of salt-bridge and cation-pi interactions, the NAMD energy plugin of VMD was used. Pore analysis of every system was carried out with MOLE2.0 (32). The last frame of every trajectory file was extracted into a pdb file using VMD. Thereafter, using these pdb files, appropriate pore profiles were obtained using MOLE2.0. A Tcl script was developed and used to track the z-position of the sodium ion to track its relative position to the DEKA ring. Dynamic sodium ion movements within the pores were modeled using VMD movie maker.

### Statistical analyses

Data from electrophysiological experiments were graphically presented using GraphPad Prism 9 software and expressed as mean ± SEM. Statistical comparisons were performed using One-Way ANOVA followed by Dunnett post-hoc tests and statistical significance was assumed when p < 0.05.

## Author contributions

ZFH, MK, SCS, TR, CLHH and APJ designed the study. ZFH performed the electrophysiological, biochemical and cell-biological experiments with help and guidance from SCS, CLHH and APJ. MK performed the molecular dynamics simulations with help and guidance from TR. APJ wrote the manuscript with help from ZFH, MK, SCS, TR and CLHH.

## Acknowledgements

ZFH was supported by an Islamic Development Bank Cambridge International Scholarship. MK was supported by funds from the University of Cambridge. SCS was supported by British Heart Foundation project grants PG/14/79/31102 and PG/19/59/34582 (to SCS, CLHH and APJ). CLLH was supported by Medical Research Council project grants MR/M001288/1 and Wellcome Trust Award 105727/Z/14/Z.

## Abbreviations

ANOVA: Analysis of variance
BrS: Brugada syndrome
Cryo-EM: cryogenic electron microscopy
DTT: Dithiothreitol
ECL: Enhanced chemiluminescence
ER: Endoplasmic reticulum
HRP: Horseradish peroxidase
MD: Molecular dynamics
PDB: Protein database
RMSD: Root-mean-square deviation of atomic positions
SND: Sinus node dysfunction

## Notes

**Conflict of interest statement:** The authors have declared that no conflict of interest exists.

### Competing Interest Statement

The authors have declared no competing interest.

## References

1. Abriel H. Cardiac sodium channel Na(v)1.5 and interacting proteins: Physiology and pathophysiology. J Mol Cell Cardiol. 2010;48(1):2–11.

2. Jiang D, Shi H, Tonggu L, Gamal El-Din TM, Lenaeus MJ, Zhao Y, et al. Structure of the Cardiac Sodium Channel. Cell. 2020;180(1):122–34 e10.

3. Oelstrom K, Goldschen-Ohm MP, Holmgren M, and Chanda B. Evolutionarily conserved intracellular gate of voltage-dependent sodium channels. Nat Commun. 2014;5:3420.

4. Ahern CA, Payandeh J, Bosmans F, and Chanda B. The hitchhiker's guide to the voltage-gated sodium channel galaxy. J Gen Physiol. 2016;147(1):1–24.

5. Favre I, Moczydlowski E, and Schild L. On the structural basis for ionic selectivity among Na+, K+, and Ca2+ in the voltage-gated sodium channel. Biophys J. 1996;71(6):3110–25.

6. Stephens RF, Guan W, Zhorov BS, and Spafford JD. Selectivity filters and cysteine-rich extracellular loops in voltage-gated sodium, calcium, and NALCN channels. Front Physiol. 2015;6:153.

7. Abriel H, and Zaklyazminskaya EV. Cardiac channelopathies: genetic and molecular mechanisms. Gene. 2013;517(1):1–11.

8. Zhang Y, Guzadhur L, Jeevaratnam K, Salvage SC, Matthews GD, Lammers WJ, et al. Arrhythmic substrate, slowed propagation and increased dispersion in conduction direction in the right ventricular outflow tract of murine Scn5a+/− hearts. Acta Physiol (Oxf). 2014;211(4):559–73.

9. Baroudi G, Napolitano C, Priori SG, Del Bufalo A, and Chahine M. Loss of function associated with novel mutations of the SCN5A gene in patients with Brugada syndrome. Can J Cardiol. 2004;20(4):425–30.

10. Veerman CC, Wilde AA, and Lodder EM. The cardiac sodium channel gene SCN5A and its gene product NaV1.5: Role in physiology and pathophysiology. Gene. 2015;573(2):177–87.

11. Zhang Y, Wang T, Ma A, Zhou X, Gui J, Wan H, et al. Correlations between clinical and physiological consequences of the novel mutation R878C in a highly conserved pore residue in the cardiac Na+ channel. Acta Physiol (Oxf). 2008;194(4):311–23.

12. Maury P, Moreau A, Hidden-Lucet F, Leenhardt A, Fressart V, Berthet M, et al. Novel SCN5A mutations in two families with "Brugada-like" ST elevation in the inferior leads and conduction disturbances. J Interv Card Electrophysiol. 2013;37(2):131–40.

13. Kapplinger JD, Tester DJ, Alders M, Benito B, Berthet M, Brugada J, et al. An international compendium of mutations in the SCN5A-encoded cardiac sodium channel in patients referred for Brugada syndrome genetic testing. Heart Rhythm. 2010;7(1):33–46.

14. Musafia B, Buchner V, and Arad D. Complex salt bridges in proteins: statistical analysis of structure and function. J Mol Biol. 1995;254(4):761–70.

15. Barlow DJ, and Thornton JM. Ion-pairs in proteins. J Mol Biol. 1983;168(4):867–85.

16. Gallivan JP, and Dougherty DA. Cation-pi interactions in structural biology. Proc Natl Acad Sci U S A. 1999;96(17):9459–64.

17. Jiang R, Martz A, Gonin S, Taly A, de Carvalho LP, and Grutter T. A putative extracellular salt bridge at the subunit interface contributes to the ion channel function of the ATP-gated P2X2 receptor. J Biol Chem. 2010;285(21):15805–15.

18. Venkatachalan SP, and Czajkowski C. A conserved salt bridge critical for GABA(A) receptor function and loop C dynamics. Proc Natl Acad Sci U S A. 2008;105(36):13604–9.

19. Zhao WS, Sun MY, Sun LF, Liu Y, Yang Y, Huang LD, et al. A Highly Conserved Salt Bridge Stabilizes the Kinked Conformation of beta2,3-Sheet Essential for Channel Function of P2X4 Receptors. J Biol Chem. 2016;291(15):7990–8003.

20. DeCaen PG, Yarov-Yarovoy V, Zhao Y, Scheuer T, and Catterall WA. Disulfide locking a sodium channel voltage sensor reveals ion pair formation during activation. Proc Natl Acad Sci U S A. 2008;105(39):15142–7.

21. Salvage SC, Zhu W, Habib ZF, Hwang SS, Irons JR, Huang CLH, et al. Gating control of the cardiac sodium channel Nav1.5 by its beta3-subunit involves distinct roles for a transmembrane glutamic acid and the extracellular domain. J Biol Chem. 2019.

22. Laedermann CJ, Syam N, Pertin M, Decosterd I, and Abriel H. beta1- and beta3-voltage-gated sodium channel subunits modulate cell surface expression and glycosylation of Nav1.7 in HEK293 cells. Front Cell Neurosci. 2013;7:137.

23. Tarradas A, Selga E, Beltran-Alvarez P, Perez-Serra A, Riuro H, Pico F, et al. A novel missense mutation, I890T, in the pore region of cardiac sodium channel causes Brugada syndrome. PLoS One. 2013;8(1):e53220.

24. Barnes S, and Hille B. Veratridine modifies open sodium channels. J Gen Physiol. 1988;91(3):421–43.

25. Denac H, Mevissen M, and Scholtysik G. Structure, function and pharmacology of voltage-gated sodium channels. Naunyn Schmiedebergs Arch Pharmacol. 2000;362(6):453–79.

26. Spencer CI. Actions of ATX-II and other gating-modifiers on Na(+) currents in HEK-293 cells expressing WT and DeltaKPQ hNa(V) 1.5 Na(+) channels. Toxicon. 2009;53(1):78–89.

27. Beu TA. Molecular dynamics simulations of ion transport through carbon nanotubes. III. Influence of the nanotube radius, solute concentration, and applied electric fields on the transport properties. J Chem Phys. 2011;135(4):044516.

28. Ahmed M, Jalily Hasani H, Ganesan A, Houghton M, and Barakat K. Modeling the human Nav1.5 sodium channel: structural and mechanistic insights of ion permeation and drug blockade. Drug Des Devel Ther. 2017;11:2301–24.

29. Humphrey W, Dalke A, and Schulten K. VMD: visual molecular dynamics. J Mol Graph. 1996;14(1):33–8, 27-8.

30. Xia M, Liu H, Li Y, Yan N, and Gong H. The mechanism of Na(+)/K(+) selectivity in mammalian voltage-gated sodium channels based on molecular dynamics simulation. Biophys J. 2013;104(11):2401–9.

31. Shen H, Liu D, Wu K, Lei J, and Yan N. Structures of human Nav1.7 channel in complex with auxiliary subunits and animal toxins. Science. 2019;363(6433):1303–8.

32. Sehnal D, Svobodova Varekova R, Berka K, Pravda L, Navratilova V, Banas P, et al. MOLE 2.0: advanced approach for analysis of biomacromolecular channels. J Cheminform. 2013;5(1):39.

33. Phillips BP, Gomez-Navarro N, and Miller EA. Protein quality control in the endoplasmic reticulum. Curr Opin Cell Biol. 2020;65:96–102.

34. Gvritishvili AG, Gribenko AV, and Makhatadze GI. Cooperativity of complex salt bridges. Protein Sci. 2008;17(7):1285–90.

35. Ing C, and Pomes R. Simulation Studies of Ion Permeation and Selectivity in Voltage-Gated Sodium Channels. Curr Top Membr. 2016;78:215–60.

36. Sun YM, Favre I, Schild L, and Moczydlowski E. On the structural basis for size-selective permeation of organic cations through the voltage-gated sodium channel. Effect of alanine mutations at the DEKA locus on selectivity, inhibition by Ca2+ and H+, and molecular sieving. J Gen Physiol. 1997;110(6):693–715.

37. Zhorov BS, and Tikhonov DB. Potassium, sodium, calcium and glutamate-gated channels: pore architecture and ligand action. J Neurochem. 2004;88(4):782–99.

38. Shen H, Li Z, Jiang Y, Pan X, Wu J, Cristofori-Armstrong B, et al. Structural basis for the modulation of voltage-gated sodium channels by animal toxins. Science. 2018;362(6412).

39. Hilber K, Sandtner W, Zarrabi T, Zebedin E, Kudlacek O, Fozzard HA, et al. Selectivity filter residues contribute unequally to pore stabilization in voltage-gated sodium channels. Biochemistry. 2005;44(42):13874–82.

40. Salvage SC, Rees JS, McStea A, Hirsch M, Wang L, Tynan CJ, et al. Supramolecular clustering of the cardiac sodium channel Nav1.5 in HEK293F cells, with and without the auxiliary beta3-subunit. FASEB J. 2020;34(3):3537–53.

41. Adrian RH. Charge movement in the membrane of striated muscle. Annu Rev Biophys Bioeng. 1978;7:85–112.

42. Sheets MF, Fozzard HA, and Hanck DA. Important Role of Asparagines in Coupling the Pore and Votage-Sensor Domain in Voltage-Gated Sodium Channels. Biophys J. 2015;109(11):2277–86.

43. Zhu W, Voelker TL, Varga Z, Schubert AR, Nerbonne JM, and Silva JR. Mechanisms of noncovalent beta subunit regulation of NaV channel gating. J Gen Physiol. 2017.

44. Pan X, Li Z, Zhou Q, Shen H, Wu K, Huang X, et al. Structure of the human voltage-gated sodium channel Nav1.4 in complex with beta1. Science. 2018;362(6412).

45. Pan X, Li Z, Huang X, Huang G, Gao S, Shen H, et al. Molecular basis for pore blockade of human Na(+) channel Nav1.2 by the mu-conotoxin KIIIA. Science. 2019;363(6433):1309–13.

46. Huang W, Liu M, Yan SF, and Yan N. Structure-based assessment of disease-related mutations in human voltage-gated sodium channels. Protein Cell. 2017;8(6):401–38.

47. Cox JJ, Sheynin J, Shorer Z, Reimann F, Nicholas AK, Zubovic L, et al. Congenital insensitivity to pain: novel SCN9A missense and in-frame deletion mutations. Hum Mutat. 2010;31(9):E1670–86.

48. Piontkivska H, and Hughes AL. Evolution of vertebrate voltage-gated ion channel alpha chains by sequential gene duplication. J Mol Evol. 2003;56(3):277–85.

49. Guo D, and Jenkinson S. Simultaneous assessment of compound activity on cardiac Nav1.5 peak and late currents in an automated patch clamp platform. J Pharmacol Toxicol Methods. 2019;99:106575.

50. Phillips JC, Braun R, Wang W, Gumbart J, Tajkhorshid E, Villa E, et al. Scalable molecular dynamics with NAMD. J Comput Chem. 2005;26(16):1781–802.

51. Feller SE, and MacKerell AD. An Improved Empirical Potential Energy Function for Molecular Simulations of Phospholipids. The Journal of Physical Chemistry B. 2000;104(31):7510–5.

52. Noskov SY, Berneche S, and Roux B. Control of ion selectivity in potassium channels by electrostatic and dynamic properties of carbonyl ligands. Nature. 2004;431(7010):830–4.

53. Masood TB, Sandhya S, Chandra N, and Natarajan V. CHEXVIS: a tool for molecular channel extraction and visualization. BMC Bioinformatics. 2015;16:119.

54. Wu J, Yan Z, Li Z, Yan C, Lu S, Dong M, et al. Structure of the voltage-gated calcium channel Cav1.1 complex. Science. 2015;350(6267):aad2395.

