## Supplemental Figures 2, 3 for "Pathological turret mutations in the cardiac sodium channel cause long-range pore disruption"

### Supplemental Figure 2

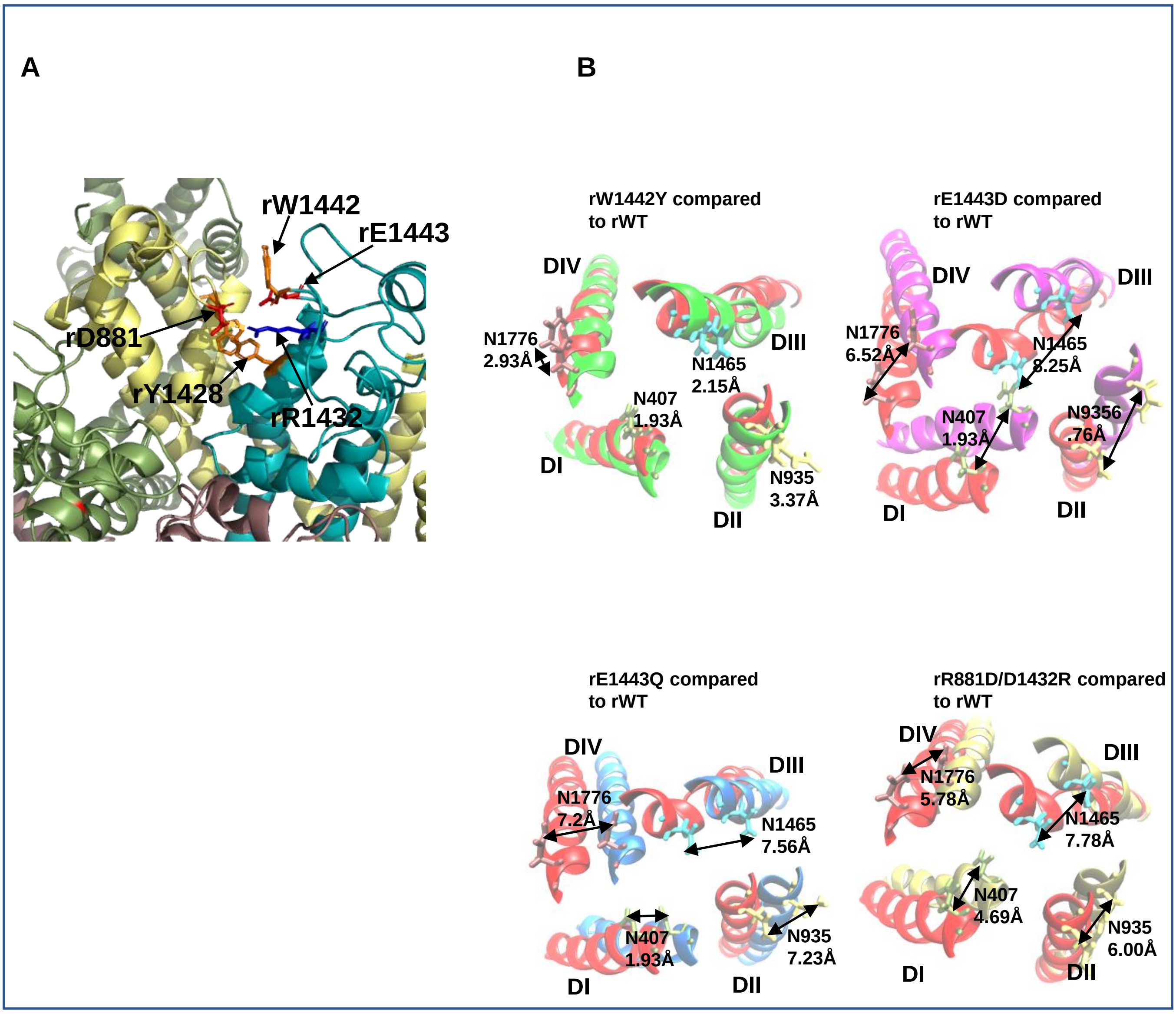


**Supplemental Figure 2. Structural perturbations in MD-simulated rat Nav1.5 mutants (A)** The final image at 15 ns of the DII-DIII interface for the simulated R881D/D1432R double mutation. Although the repositioned R1432 residue can still form a salt-bridge with D881, it can no longer form a salt-bridge with E1443. This abolishes the normal cation-π bond with W1442. **(B)** The final image at 15 ns of the S6 helices for simulated wild type (WT) and mutant rat Nav1.5 channels. The conformation of the S6 helices for each mutant is superimposed on the S6 helices for wild type Nav1.5 (red). The helices are viewed from the channel cytoplasmic face. The positions of the conserved asparagine residues noted in the text and the displacement in their positions between the mutant and wild type S6 helices are as indicated in Å.

### Supplemental Figure 3A

**A**

```
Nav1.1  GMQLFGKSYKDCV----CKIAS--DCQLPRWHMNDFFHSFLIVFRVLCGEWIETMWDCME  961
Nav1.2  GMQLFGKSYKECV----CKISN--DCELPRWHMHDFFHSFLIVFRVLCGEWIETMWDCME  952
Nav1.3  GMQLFGKSYKECV----CKIND--DCTLPRWHMNDFFHSFLIVFRVLCGEWIETMWDCME  953
Nav1.4  GMQLFGKSYKECV----CKIAL--DCNLPRWHMHDFFHSFLIVFRILCGEWIETMWDCME  771
Nav1.5  GMQLFGKNYSELR------DSD--SGLLPRWHMMDFFHAFLIIFRILCGEWIETMWDCME  908
Nav1.6  GMQLFGKSYKECV----CKINQ--DCELPRWHMHDFFHSFLIVFRVLCGEWIETMWDCME  946
Nav1.7  GMQLFGKSYKECV----CKIND--DCTLPRWHMNDFFHSFLIVFRVLCGEWIETMWDCME  937
Nav1.8  GKQLLGENYRNNR----KNISAP-HEDWPRWHMHDFFHSFLIVFRILCGEWIENMWACME  859
Nav1.9  GMQLFGRSFNSQKSPKLCNPTGPTVSCLRHWHMGDFWHSFLVVFRILCGEWIENMWECMQ  778
        * **:*..: .                  :*** **:*:**::**:*******.** **:

Nav1.1  QVATFKGWMDIMYAAVDSRNVELQPKYEESLYMYLYFVIFIIFGSFFTLNLFIGVIIDNF  1486
Nav1.2  QVATFKGWMDIMYAAVDSRNVELQPKYEDNLYMYLYFVIFIIFGSFFTLNLFIGVIIDNF  1476
Nav1.3  QVATFKGWMDIMYAAVDSRDVKLQPVYEENLYMYLYFVIFIIFGSFFTLNLFIGVIIDNF  1471
Nav1.4  QVATFKGWMDIMYAAVDSREKEEQPQYEVNLYMYLYFVIFIIFGSFFTLNLFIGVIIDNF  1298
Nav1.5  QVATFKGWMDIMYAAVDSRGYEEQPQWEYNLYMYIYFVIFIIFGSFFTLNLFIGVIIDNF  1473
Nav1.6  QVATFKGWMDIMYAAVDSRKPDEQPKYEDNIYMYIYFVIFIIFGSFFTLNLFIGVIIDNF  1467
Nav1.7  QVATFKGWTIIMYAAVDSVNVDKQPKYEYSLYMYIYFVVFIIFGSFFTLNLFIGVIIDNF  1460
Nav1.8  QVATFKGWMDIMYAAVDSREVNMQPKWEDNVYMYLYFVIFIIFGGFFTLNLFVGVIIDNF  1421
Nav1.9  QVATFKGWMDIIYAAVDSTEKEQQPEFESNSLGYIYFVVFIIFGSFFTLNLFIGVIIDNF  1311
        ********  *:******   . ** :* .   *:***:*****.*******:*******
```

* Sites that are fully conserved between all channels.

: Sites that show conservative changes between channels

. Sites that show semi-conservative changes between channels

**Supplemental Figure 3. Conservation of the DII-DIII turret interface (A)** Sequence alignments of human Nav 1.5 (bold), compared to other human Nav channel isoforms. The conservation of the DII-DIII turret interface residues are indicated. Residues in Nav1.1 that are mutated in some forms of inherited epileptic encephalopathy (46) and the residue in Nav1.7 (R907) that is mutated in a case of congenital insensitivity to pain (47) are bordered by a black square.

### Supplemental Figure 3B and C

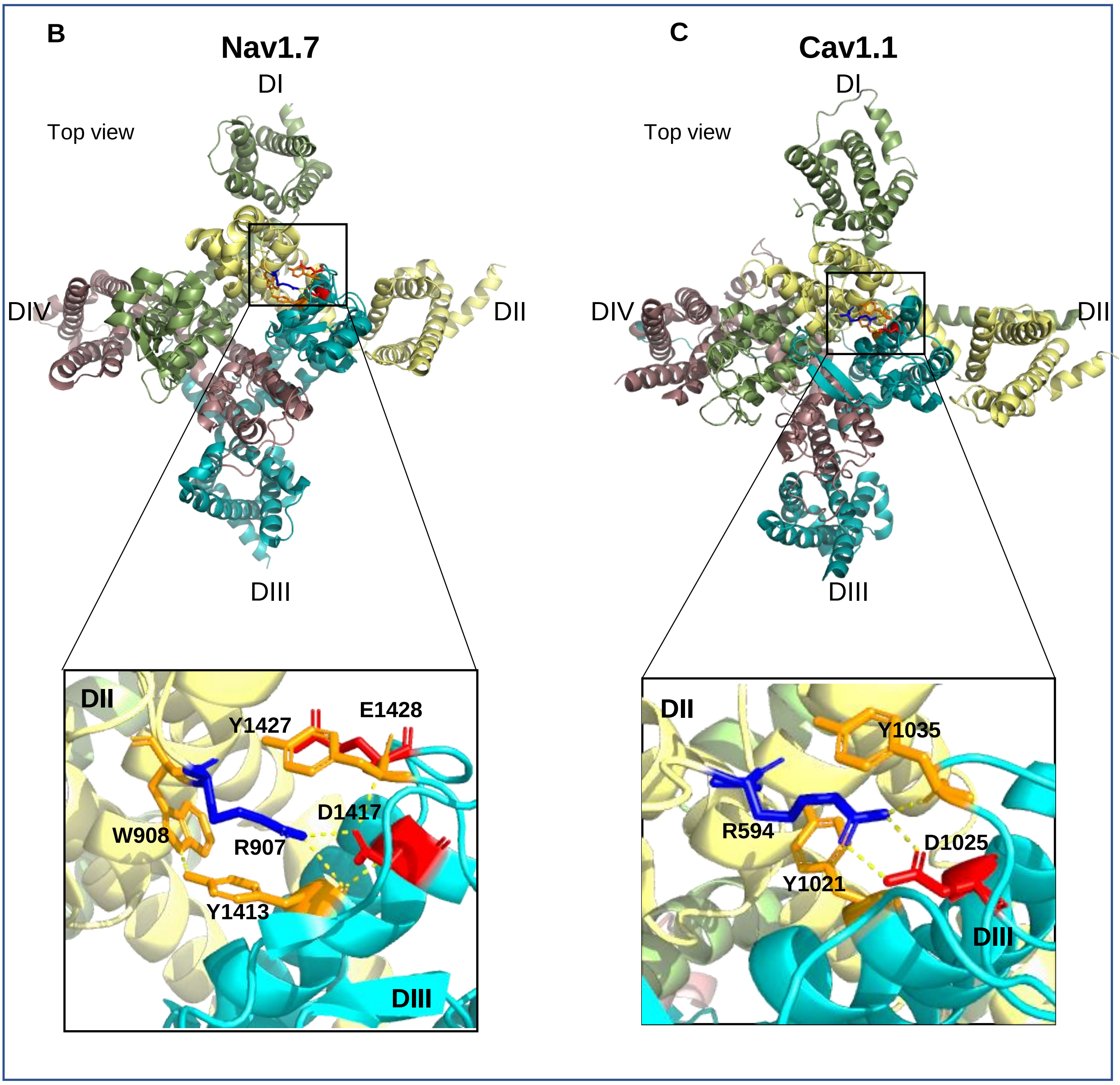


**Supplemental Figure 3. Conservation of the DII-DIII turret interface (B)** A cartoon representation of human $Na_v$1.7 (PDB ID: 6j8j), showing the conserved DII-DIII turret interface residues, including the R907 residue noted above. **(C)** A cartoon representation of rabbit Cav1.1 (PDB ID: 5GJV) (54), showing a similar salt-bridge and cation-π bonded DII-DIII turret interface, with relevant amino acid residues indicated.
